# BAG2 Condensates Couple Proteostasis to CD8^+^T Cell Surveillance

**DOI:** 10.64898/2026.04.24.719751

**Authors:** Maria C. Almeida, Tian Wang, Andrew P. Longhini, Samuel Lobo, Carolina M. Camargo, Eric D. Tinkle, Mookwang Kwon, Guilherme Z. Duarte, Isabel O. Hirsch, Cesar A.J. Ribeiro, Fernando A. Oliveira, M. Scott Shell, Joan-Emma Shea, Judith A. Steen, Kenneth S. Kosik, Daniel C. Carrettiero

## Abstract

Protein aggregation, impaired degradation, and immune activation are central hallmarks of neurodegenerative diseases, yet how these processes are coordinated remains unclear. Here, we identify Immune-Protein Degradation Bodies (I-PDBs), a previously unrecognized class of BAG2-driven, phase-separated organelles that integrate protein quality control with adaptive immunity. IFNγ induce I-PDB formation at the endoplasmic reticulum (ER), where they concentrate immunoproteasome components, MHC-I peptide-loading machinery, and ER-associated chaperones. I-PDBs redirect proteostatic cargo from centrosomal aggregation pathways to spatially restricted degradation sites optimized for antigenic peptide generation, coupling selective substrate clearance to CD8⁺ T cell engagement. Using a cellular model of aggregation-prone tau, we show that I-PDBs capture pathological tau fibrils at ER–microtubule interfaces and process them into potentially antigenic peptides, thus reducing the load of aggregation-prone tau peptides. We term this mechanism the Proteostasis-Associated Immune Relay (PAIR), establishing I-PDBs as critical hubs linking proteostasis to immune surveillance with broad implications for disease.

**Graphical Abstract:** 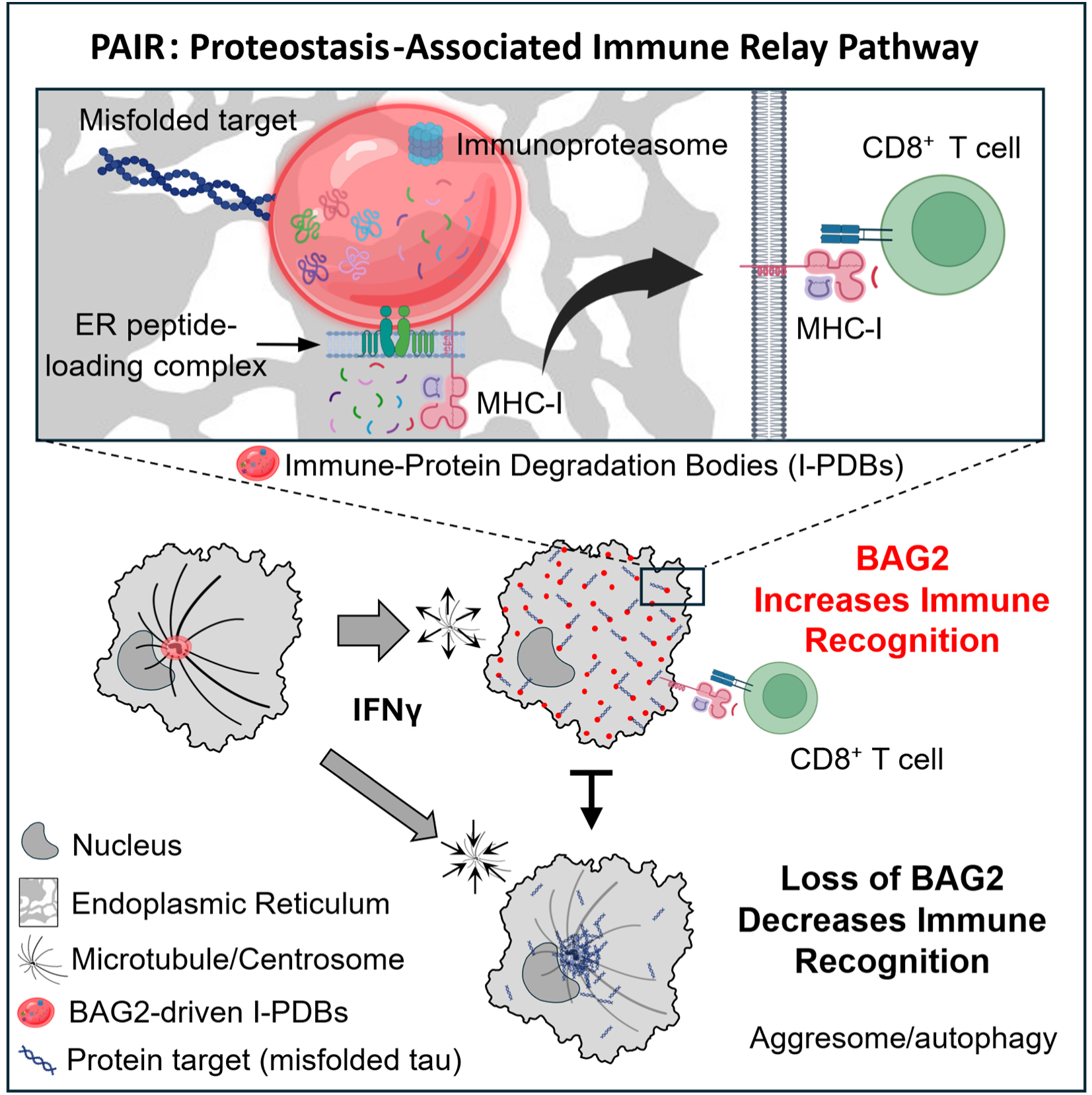

**Highlights:**

- IFNγ drives BAG2-dependent Immune-Protein Degradation Bodies (I-PDBs)
- I-PDBs assemble at the endoplasmic reticulum and are enriched in immunoproteasome and MHC-I machinery
- I-PDBs shunt misfolded proteins, including pathological tau, away from aggresomes
- I-PDBs couple proteostasis to antigen presentation, enhancing CD8⁺ T cell recognition
- The Proteostasis-Associated Immune Relay (PAIR) defines a pathway linking proteostasis to adaptive immunity

## INTRODUCTION

Proteome integrity relies not only on intracellular protein quality control but also on immune surveillance mechanisms that determine outcomes from cell survival to cell elimination^1,2^. While the pathways governing intracellular proteostasis are well-characterized, how they intersect with the immune system remains unclear. This question is particularly pressing in neurodegenerative disorders such as Alzheimer’s disease, which are characterized by chronic inflammation and the accumulation of misfolded or aggregated proteins^3–7^.

Biomolecular condensates formed through liquid–liquid phase separation (LLPS) have emerged as fundamental organizers of intracellular biochemistry^8,9^. In the context of protein degradation, LLPS can organize protein quality control pathways, coordinating proteasomal degradation with autophagic clearance under proteotoxic stress or substrate overload^10–12^. This spatial organization ensures efficient substrate processing and allows the cell to respond dynamically to stress and protein aggregation. BAG2 (Bcl-2-associated athanogene 2), a co-chaperone that targets aggregation-prone substrates such as tau, undergoes LLPS capable of organizing protein quality control via the ubiquitin-independent proteasome pathway^13,14^. Substrates that are not efficiently processed along this route can accumulate in aggresome-like structures and be redirected to alternative proteostasis pathways.

Beyond intrinsic proteostasis, immune surveillance also provides a systemic layer of quality control. CD8⁺ cytotoxic T-lymphocytes monitor cellular integrity by sensing peptides on the cell surface displayed on major histocompatibility complex class I (MHC-I) molecules^15–17^. These peptides, derived from misfolded, aggregated, mutated, or foreign proteins, are largely generated by proteasome degradation^15,16,18–20^ and trimmed in the endoplasmic reticulum (ER) to 8–10 amino acids^22,23^ where they are loaded into MHC-I molecule. Interferon-γ (IFNγ), a cytokine central to cell-mediated immunity and released by CD8⁺ T cells, induces a pro-inflammatory state, including induction of the immunoproteasome^15^ which enhances peptide generation by over 100-fold^25^. Three constitutive catalytic subunits of the 20S proteasome, PSMB6 (β1), PSMB7 (β2), and PSMB5 (β5) are replaced by catalytic subunits β1i (PSMB9), β2i (PSMB10), and β5i (PSMB8) forming the immunoproteasome. However, antigen presentation is intrinsically inefficient, with only a small fraction (∼1 peptide : 10,000 proteins degraded) ultimately sampled for MHC-I presentation^24^, raising the question of how cells select antigenic peptides. While this remodeling increases immune surveillance capacity, it simultaneously raises the burden of proteolytic products. How cells coordinate increased peptide production with selective antigen presentation remains unresolved, particularly for aggregation-prone substrates.

Tauopathies are characterized by the intracellular accumulation of fibrillar tau, which is closely associated with neuronal dysfunction and degeneration^26^. Tau aggregation primarily originates in neurons, and can propagate through seeding mechanisms^27^. Extracellular tau is captured by glia and microglia and drained via meningeal lymphatics^28,29^, extending the impact of tau pathology beyond the CNS^30^. Blocking lymphatics flux leads to tau and CD8⁺T cell accumulation inside the brain^31^. Additional evidence for the interplay between the periphery and related tauopathies comes from observations that IFNγ signaling and clonally expanded CD8⁺ T cells are increased in affected brain regions^32–36^. In parallel, brain-derived antigens such as tau can be presented in secondary lymphoid tissues, where they may prime CD8⁺ T cell^37^. Recent studies further indicate that tau-driven proteostatic stress elicits a coordinated innate and adaptive immune response characterized by microglial activation and infiltration of cytotoxic CD8⁺ T cells, whose abundance correlates with neuronal loss^38^.

We identify a new class of BAG2-driven, phase-separated organelle, the Immune-Protein Degradation Bodies (I-PDBs), that provide a local environment for protein degradation and the generation of antigenic peptides. IFNγ drive the formation of I-PDBs, which are enriched in immunoproteasome components, MHC-I loading machinery, and chaperones closely associated with the endoplasmic reticulum (ER). This reorganization shifts proteolytic activity away from centrosomal aggregation routes toward immune-specialized sites, couples substrate capture to peptide generation, and ensures the production of optimally sized antigenic fragments, thereby enhancing CD8⁺ T cell engagement. I-PDBs capture pathological tau fibrils at ER–microtubule interfaces and process them into antigenic peptides, reducing tau peptides that contain the aggregation-prone PHF6 motif. Collectively, these findings reveal a cellular strategy for translating intracellular stress into immune signals, which we term the Proteostasis-Associated Immune Relay (PAIR) pathway, establishing I-PDBs as critical hubs with potential implications for multiple disease states.

## RESULTS

### IFNγ induces immune-protein degradation bodies (I-PDBs) and tunes antigen processing

BAG2 forms sucrose-induced degradation condensates, which have been previously characterized by our group^13^. Here, we show that IFNγ, a cytokine elevated in tauopathies^32–35^, also induces BAG2 condensates in an inflammatory context. Upon IFNγ (50 ng/ml, 24 h), BAG2 transitioned from a diffuse cytoplasmic distribution to dynamic phase-separated condensates in H4 cells, as observed by live-cell imaging of Clover-BAG2–expressing cells (Figure 1A) and by immunofluorescence detecting endogenous BAG2 (Figure S1A). IFNγ increased BAG2 expression as confirmed by immunoblotting (Figure 1B) and qPCR (Figure S1B). Consistent with its canonical effects, IFNγ upregulated MHC-I and the immunoproteasome subunit PSMB8, with MHC-I redistributing to the cell periphery, suggesting enrichment at the plasma membrane (Figure S1C-E). BAG2 knockdown (KD), previously established^13^ and validated here (Figure S1F), did not affect MHC-I levels, IFNγ-induced MHC-I upregulation or MHC-I peripheral redistribution (Figure S1D,E). With IFNγ treatment, BAG2 condensates recruited PSMB8, as shown by immunofluorescence (Figure 1C). Co-transfection of H4 cells with Clover-BAG2 and Ruby-PSMB8 confirmed this recruitment and revealed partial dynamic demixing, with condensates undergoing fusion and fission (Figure 1D; Video S1). We hereafter refer to these IFNγ–induced, immunoproteasome-enriched BAG2 condensates as Immune-Protein Degradation Bodies (I-PDBs).

**Figure 1.**
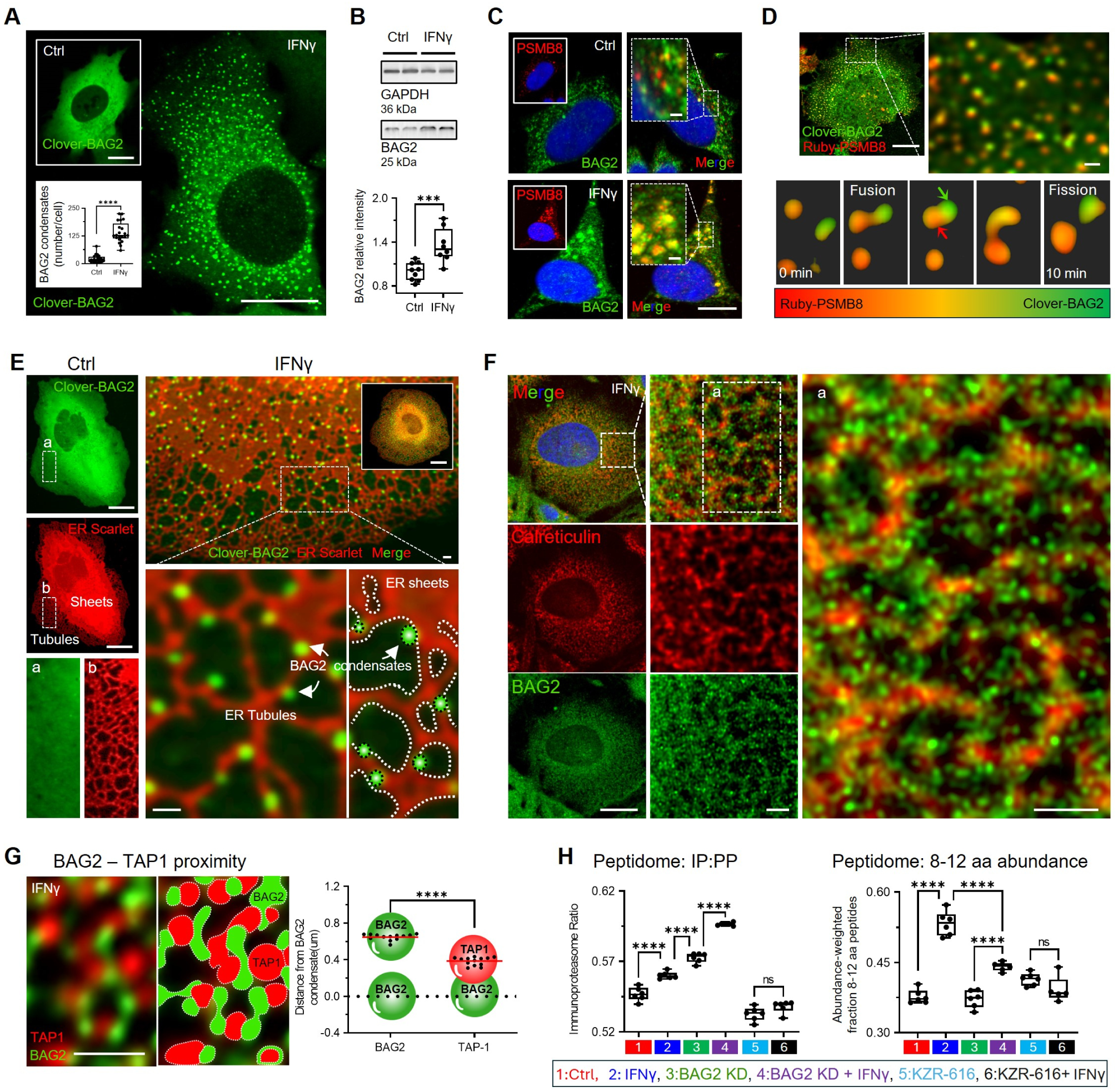
IFNγ induces immune-protein degradation bodies (I-PDBs) and tunes antigen processing. (A) Live-cell imaging of Clover-BAG2–expressing H4 cells in untreated or IFNγ–treated cells (50 ng/mL, 24 h), with quantification of condensates per cell. Dots represent mean values per field of view (FOV) from independent replicates (n=21-22, with 9-57 cells/FOV). (B) BAG2 protein (immunoblot) in untreated and IFNγ–treated cells. (C) Immunofluorescence of BAG2 and PSMB8 in untreated and IFNγ–treated cells. (D) Live-cell imaging of cells co-expressing Clover-BAG2 and Ruby-PSMB8, showing dynamic interactions. Arrows indicate BAG2 (green) and PSMB8 (red) enrichment. (E) Live-cell imaging of H4 cells co-expressing ER-Scarlet and Clover-BAG2, showing stimulus-dependent redistribution of BAG2 condensates relative to the ER following IFNγ (50 ng/mL, 24 h). (F) Immunofluorescence staining of calreticulin and BAG2. (G) Immunofluorescence of BAG2 and TAP1 with quantification of spatial proximity. Plot shows nearest-neighbor distances between BAG2 condensates and between BAG2 and TAP1 signals. Each dot represents the mean of one FOV from independent replicates (n=13 per condition, with 800-2000 BAG2 condensates/FOV). (H) Immunoproteasome contribution to peptide generation across indicated conditions. All box plots show interquartile range, median, and min/max (whiskers); (B,H) Dots represent independent replicates; (A, B) two-tailed t test; (G) two-way ANOVA with Tukey’s test (***p < 0.001; ****p < 0.0001; ns). Scale bars: 10 µm and 2 µm (zoom). See also Figure S1, Table S1 and Videos S1-2.

Given that immunoproteasome activity supplies peptides for ER-based antigen processing, we asked whether I-PDBs physically associate with the ER. We tested this by co-expressing ER–mScarlet and Clover–BAG2 in H4 cells under IFNγ. I-PDBs localized along the outer ER membrane of an interconnected ER network of sheets and tubules spanning the cytoplasm (Figure 1E, Video S2). This spatial association was confirmed using endogenous immunostaining for BAG2 and the ER marker Calreticulin, which revealed small BAG2 puncta closely apposed to the ER under IFNγ (Figure 1F).

Consistent with a role in antigen processing, immunofluorescence showed that I-PDBs were positioned near TAP1 transporter clusters (Figure 1G), the ER-resident channel responsible for delivering immunoproteasome-generated peptides for MHC-I loading. Quantitative analysis revealed that the distance between I-PDBs and TAP1 is significantly shorter than between I-PDBs, indicating that I-PDBs preferentially localize near the TAP1 clusters rather than being randomly distributed throughout the cytoplasm.

To test if I-PDBs influence antigen processing, we performed peptidomics under six conditions: untreated, IFNγ, BAG2 KD, BAG2 KD + IFNγ, the immunoproteasome inhibitor KZR-616, and KZR-616 + IFNγ. We first evaluated the immunoproteasome (IP)-to-proteasome (PP) peptide ratio (IP:PP), classifying peptides by their C-terminal residues (Methods). As expected, IFNγ increased the IP:PP ratio (Figure 1H – left panel). BAG2 KD alone also elevated the IP:PP ratio, with IFNγ producing an additional increase. KZR-616 decreased the IP:PP ratio in control cells and abolished the IFNγ effect, confirming immunoproteasome dependence. Total peptide production showed a similar pattern (Figure S1G), indicating that BAG2 limits the relative contribution of immunoproteasome-derived peptides. We next evaluated the IP enrichment peptides within the MHC-I length range (8–12 aa). IFNγ increased the 8–12 aa peptide fraction (Figure 1H – right panel). BAG2 KD did not increase this fraction, and markedly attenuated IFNγ-induced enrichment of peptides in the 8-12 aa length range. KZR-616 abolished the IFNγ effect. Together, these results indicate that BAG2 is required for efficient generation of immunoproteasome-derived peptides within the optimal MHC-I length range.

Overall, these results show that IFNγ drives the formation of dynamic I-PDBs, spatially reorganizing the local degradation machinery and tuning immunopeptide production. I-PDBs occupy a distinct proteostatic niche, positioned near the ER and TAP1 transporters, suggesting a functional interface with antigen-processing pathways. In the next section, we expand the analysis of I-PDB-associated proteins by comparing them with sucrose-induced BAG2 degradation condensates^13^, a non-inflammatory stress state, to better define I-PDBs functional role.

### IFNγ reprograms BAG2 degradation condensates into an immune state

To define the core composition of I-PDBs and identify immune-specific features distinct from stress-induced BAG2 condensation^13^ (hereafter termed S-PDBs), we performed APEX2 proximity labeling proteomics^39,40^ using BAG2 protein as bait (Figure S2A-D). H₂O₂-triggered APEX2-BAG2 biotinylated proteins within ∼20 nm (Figure S2A), allowing comparison of the APEX-BAG2 proteome under three conditions: untreated, IFNγ, and sucrose-induced osmotic stress (125 mM, 15 min), a well-characterized stimulus that rapidly induces S-PDB formation without inflammation^13^.

APEX2–BAG2 profiling identified 403 proteins in IFNγ–treated and 91 in sucrose-treated cells (Figure 2A, Table S1). Of these, 48 were shared, with 355 IFNγ-specific and 43 sucrose-specific, highlighting immune-selective recruitment to I-PDBs. When compared to untreated, IFNγ induced enrichment of immunoproteasome subunits (PSMB8, PSMB9, PSMB10), the MHC-I peptide-loading complex (HLA-A/C, B2M, TAP1, TAP2, TAPBP, ERAP1) and ER-associated folding machinery, including chaperones (CALR, CANX, HSPA5/BiP) and PDIA-family oxidoreductases (P4HB/PDIA1, PDIA3, PDIA6) (Figure 2B; Table S1), indicating BAG2 proximity to immunoproteasome machinery, peptide-loading factors and ER molecules. Enrichment pathway analysis also revealed strong associations with immune processes and proteasome-mediated protein degradation (Figure 2C). Comparison of the APEX2–BAG2 dataset with the IFNγ-treated whole-cell proteome (Methods) further confirmed its specificity (Figure S2E).

**Figure 2.**
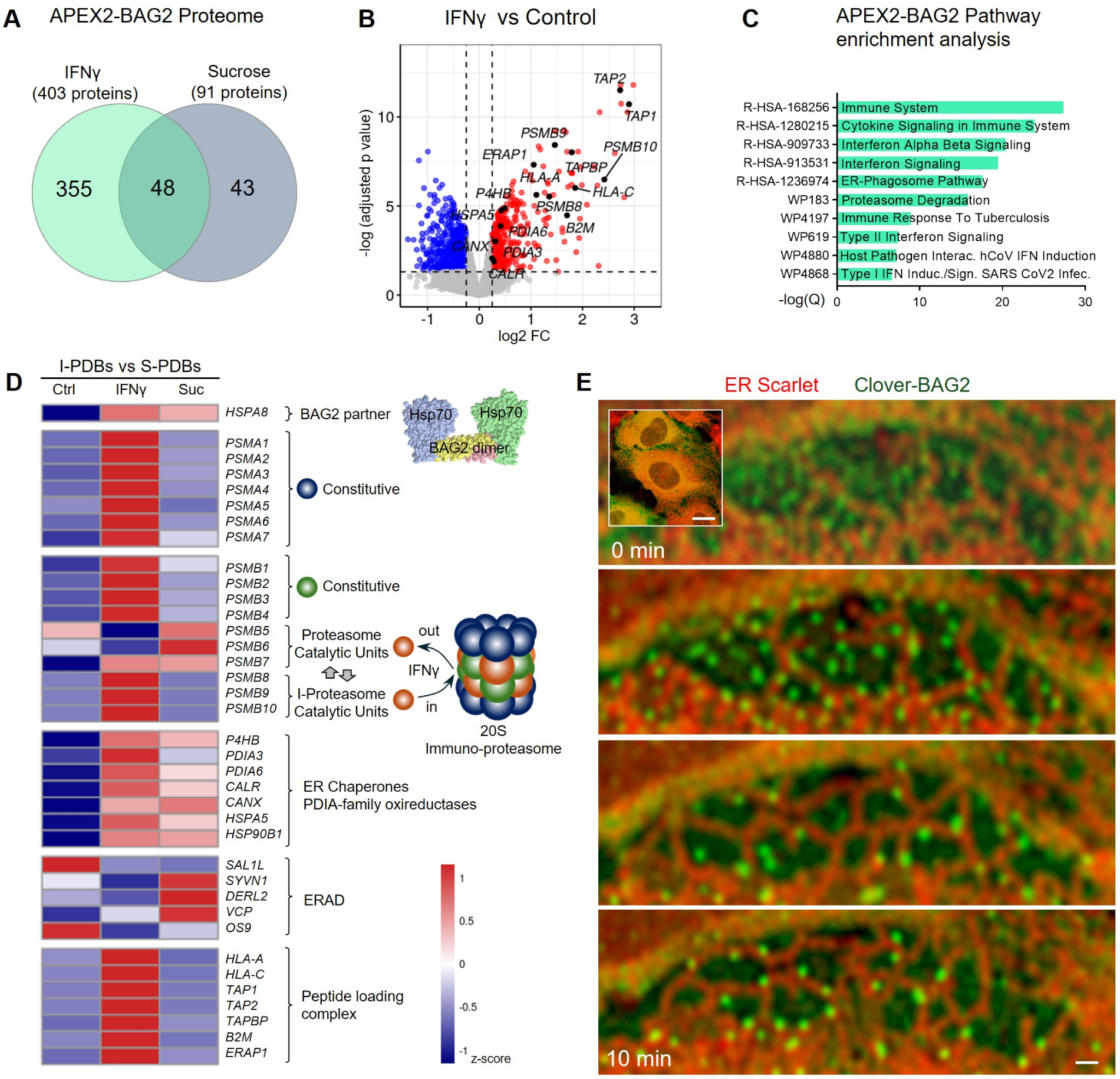
IFNγ reprograms BAG2 degradation condensates into an immune state. (A) Overlap between proteins in the APEX2-BAG2 proteome following sucrose-induced osmotic stress or IFNγ treatment, revealing stimulus-specific remodeling. (B) Volcano plot of proteins enriched in IFNγ–induced BAG2 condensates versus controls (gene symbols shown). (C) Pathway enrichment of BAG2-associated proteins (Reactome, WikiPathways). (D) Heatmap of proteasome, immunoproteasome, ER chaperone, and ERAD-associated proteins in the APEX2-BAG2 proximal proteome of untreated, IFNγ–treated, and sucrose (Suc)-treated cells (z-score; mean normalized abundance, n=4 per condition). Gene symbols shown. (E) Time-lapse imaging of BAG2 condensate formation and remodeling during osmotic stress (sucrose, 125 mM). Scale bar: 10 µm and 2 µm (zoom). See also Figures S2-3, Table S1 and Video S3

Having established the core composition of the I-PDBs, we next compared the recruitment of key functional modules in I-PDBs to S-PDBs relative to control (Figure 2D). HSP70, BAG2’s primary molecular partner in client processing^27^, was enriched in both I-PDBs and S-PDBs. The E3 ubiquitin ligase CHIP (STUB1), free ubiquitin (UBB, UBC) and ubiquitin–ribosomal fusion proteins (UBA52, RPS27A) were not detected (Table S1), and practically all 19S proteasome regulatory subunits, including base ATPases (PSMC1–6) and lid components (PSMD1–PSMD14) were absent in both condensates (Table S1) consistent with the function of BAG2 condensates in ubiquitin-independent degradation^13,14^. Strikingly, the complete ISGylation machinery^41^ were found in I-PDBs but not in S-PDBs, including UBA7 (E1), UBE2L6 (E2), HERC6 (E3), ISG15, USP18, DTX3L, and the ISG15 adaptor NUB1 (Figure S2F).

Consistent with this model, multiple 20S proteasome constitutive core subunits (PSMA1–7 and PSMB1–4) were enriched in I-PDBs but not in S-PDBs (Figure 2D). Unlike S-PDBs, where proteasome representation was limited and largely restricted to catalytic β subunits (PSMB5–7), I-PDBs showed broader recruitment, including constitutive (PSMA1–7 and PSMB1–4) and immunoproteasome (PSMB8–10) components, consistent with assembly of a functional 20S core.

To determine whether ER association partners are specific to I-PDBs, we compared ER-related proteins between I-PDBs and S-PDBs relative to control (Figure 2D). Both condensate types were significantly enriched in ER chaperones calnexin and BiP (CANX, HSPA5) as well as PDIA-family oxidoreductases (PDIA1, PDIA6), indicating that ER association is an intrinsic feature of both I-PDBs and S-PDBs.

The core ER-associated degradation (ERAD) components (SEL1L, SYVN1/HRD1, DERL1/2, VCP, UBE2J1, and OS9) were not enriched in I-PDBs (Figure 2D). Although canonical unfolded protein response (UPR) regulators were not detected by APEX2-BAG2 proximity labeling—and are not expected to be captured reliably due to their nuclear or distal cytosolic localization—these signaling components (XBP1/XBP1s, ATF4, CHOP, and ATF6) were also undetectable in the whole cell proteomics analyses (Methods) of IFNγ-treated cells (Table S1).

Together, these findings indicate that both I-PDBs and S-PDBs constitutively engage ER folding and entry machinery without triggering ERAD or UPR. However, only I-PDBs recruits immunoproteasome subunits (PSMB8–10), key components of the MHC-I peptide-loading machinery, including HLA-A, HLA-C, TAP1/2, TAPBP, and ERAP1, which trims peptides for optimal MHC-I loading (Figure 2D). Key differences between I-PDBs and S-PDBs are summarized in Figure S3.

In parallel, we trained a classifier to distinguish proteins enriched in the APEX2-BAG2 IFNγ proteome from those detected in the IFNγ whole-cell proteome, using ESM3 (Evolutionary Scale Modeling v3) protein language model embeddings derived from AlphaFold-predicted structural features (Methods). This model identified discriminative sequence features associated with proteins recruited to I-PDBs, achieving a median ROC AUC of 0.836. At the local sequence level, enrichment of acidic residues emerged as a characteristic sequence signature of I-PDB-associated proteins (Figure S3B,C).

To determine whether I-PDBs or S-PDBs nucleate directly at the ER or instead form in the cytoplasm and subsequently traffic to ER membranes, we used sucrose as an acute stressor for S-PDBs formation. Live-cell imaging under sucrose treatment revealed that BAG2 condensates initially emerge in the cytoplasm and then rapidly migrate toward the ER (Figure 2E; Video S3), where they establish stable and extensive contacts along ER tubules.

Collectively, these data support a model in which I-PDBs associated with the ER are widely distributed throughout the cytoplasm and function as specialized motile antigen-processing degradation hubs. In contrast, unprocessed proteins have been reported to accumulate in aggresome-like structures^13,42^ near the centrosome. This spatial segregation suggests a coordinated division of labor, with I-PDBs handling local degradation and peptide generation, and centrosome-associated pathways managing the remaining aggregation-prone material (bulk degradation), as discussed next.

### BAG2 restrains pericentrosomal trafficking to promote antigen processing

To define BAG2’s spatial organization under basal conditions, we examined its subcellular distribution by immunofluorescence. While largely diffuse in the cytoplasm, BAG2 consistently formed one or two prominent perinuclear condensates (∼0.5 µm) (Figure 3A). Co-staining with γ-tubulin confirmed their centrosomal localization (Figure 3B), which also colocalize with PSMB8, revealing a pre-existing spatial link between BAG2 condensates and basal immune-related proteostasis machinery.

**Figure 3.**
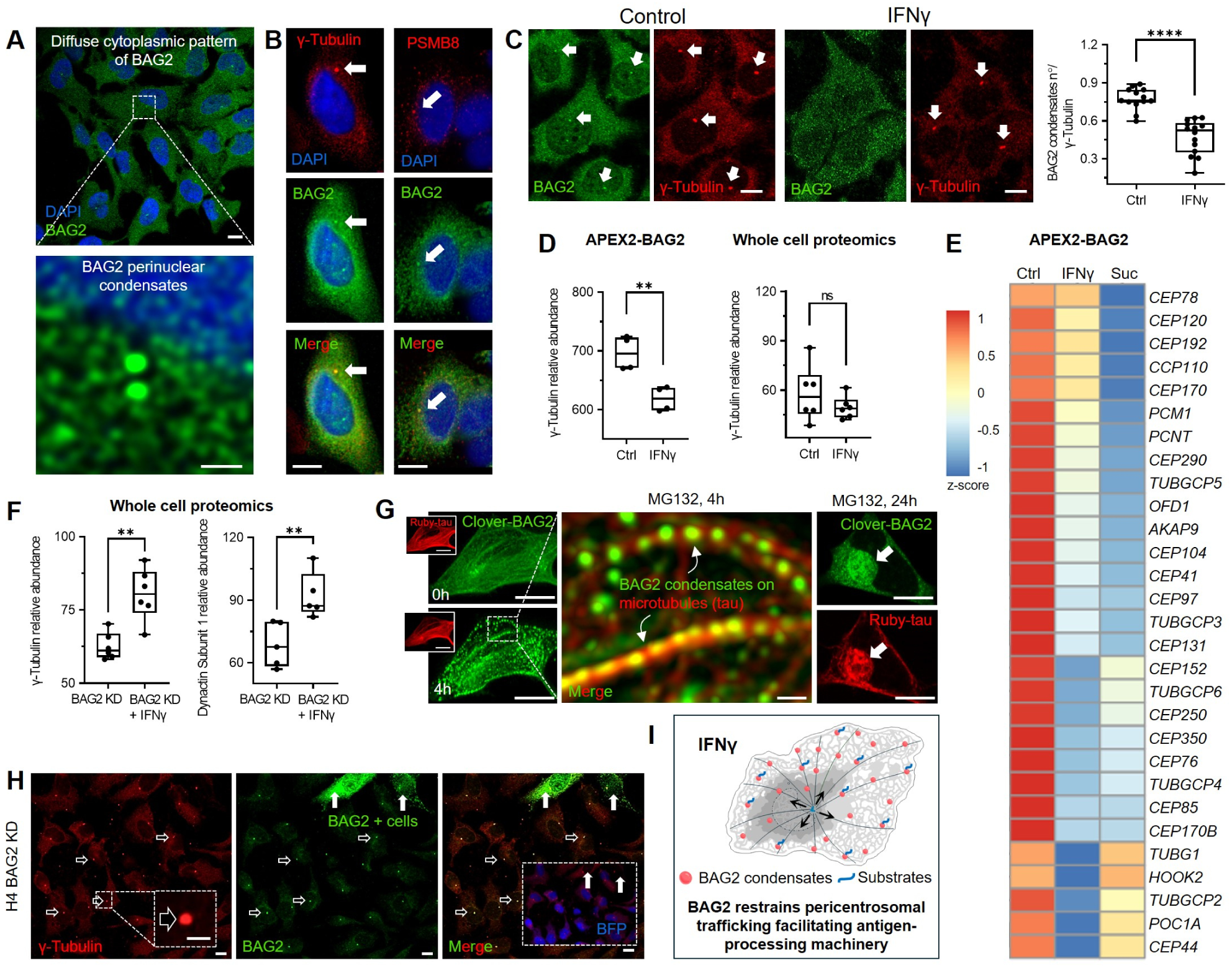
BAG2 restrains pericentrosomal trafficking to promote antigen processing. (A) Immunofluorescence of endogenous BAG2 in untreated H4 cells shows diffuse cytoplasmic distribution with perinuclear enrichment. Scale bars, 10 µm and 1 µm (zoom). (B) Co-staining of BAG2 with γ-tubulin or PSMB8 shows centrosomal localization under basal conditions. (C) Co-staining of BAG2 and γ-tubulin in untreated and IFNγ–treated cells. IFNγ reduces centrosomal BAG2, quantified as condensates per centrosome. Each dot represents one field of view (FOV) from independent replicates. An average of 60-66 cells per FOV were included in each condition. (D) γ-tubulin abundance in the APEX2-BAG2 proximal proteome decreases upon IFNγ treatment (left), while total levels remain unchanged (right). (E) Heatmap of centrosome-associated proteins in the APEX2-BAG2 proximal proteome from untreated, IFNγ–treated, and sucrose (Suc)-treated cells (z-score; mean normalized abundance, n=4 per condition). Gene symbols shown. (F) γ-tubulin and dynactin subunit 1 abundance in whole-cell proteomes of untreated and IFNγ–treated BAG2-KD cells. (G) Live-cell imaging of Ruby-tau–expressing H4 cells transiently expressing Clover-BAG2 and treated with MG132. BAG2 localizes along microtubules at 4h and accumulates with tau perinuclearly at 24 h. (H) Immunofluorescence of BAG2 and γ-tubulin in BAG2 KD cells shows residual BAG2 enrichment at centrosomes. (I) Model of BAG2-mediated pericentrosomal trafficking. Box plots show interquartile range (25th–75th percentiles), median (center line), and min–max (whiskers). Dots represent independent biological replicates. (C, D, F) two-tailed Student’s t test; **p < 0.01; ****p < 0.0001; ns, not significant. Scale bars 10 µm, 1 µm (zoom, A), 2 µm (zoom, G). See also Figure S4 and Table S1.

Upon IFNγ, BAG2 colocalization with the centrosome was no longer observed (Figure 3C), indicating redistribution, likely toward ER-associated compartments. Consistently, APEX2–BAG2 proteomics showed decreased γ-tubulin enrichment under IFNγ, while whole-cell proteomics revealed unchanged total γ-tubulin levels (Figure 3D), suggesting IFNγ-dependent relocalization of BAG2 rather than altered γ-tubulin abundance. IFNγ also reduced enrichment of multiple centrosome-associated proteins in the APEX2–BAG2 proteome (Figure 3E), a pattern similarly observed under sucrose stress, indicating that BAG2 dissociation from centrosome is not specific to inflammation but common to both I-PDBs and S-PDBs formation.

In BAG2 KD, whole-cell proteomics showed that IFNγ increased γ-tubulin (Figure 3F), reflecting expansion of the centrosome, as well as DCTN1, a dynactin subunit involved in dynein-dependent retrograde transport toward the centrosome. This expansion coincided with coordinated upregulation of centrosomal trafficking components in the absence of BAG2, including dynein heavy chain DYNC1H1, γ-tubulin complex protein TUBGCP2, and elements of the dynein–dynactin transport machinery such as the dynactin subunit ACTR1A and cargo adaptor HOOK3 (Figure S4A), reflecting enhanced microtubule-organizing capacity and dynein-dependent retrograde transport upon loss of BAG2.

Independent evidence supports BAG2’s role in stress-induced spatial trafficking. We previously showed that proteasome inhibition induces BAG2 condensation within 4h and, upon prolonged treatment (24 h), promotes BAG2 and tau co-accumulation in pericentrosomal aggresome-like structures^13^. Here, we extend these findings by resolving the spatial organization of BAG2 condensates during this process. Live-cell imaging of Clover–BAG2 and Ruby–tau, a microtubule-associated structural marker, revealed that short-term MG132 treatment (4 h, 10 µM) induces rapid BAG2 condensation along microtubule-defined trajectories (Figure 3G), preceding pericentrosomal accumulation. At later time points (24 h), BAG2 and tau co-localize within aggresome-like structures, consistent with microtubule-dependent trafficking toward the centrosome^13^.

These observations establish the aggresome as a deposition site for unresolved tau substrates under sustained proteostasis impairment. Remarkably, even in BAG2 KD cells, a residual perinuclear BAG2 signal persisted at the centrosome (Figure 3H) despite loss of diffuse cytoplasmic staining, revealing a stable centrosomal pool preferentially retained upon BAG2 depletion.

These findings support a model in which a subset of BAG2 condensates localize to the centrosome to maintain proteostasis but, upon IFNγ stimulation and I-PDB formation, redistribute to ER-associated, immune-specialized compartments with a remodeled proteome and reduced pericentrosomal trafficking (Figure 3I). How this reorganization affects handling of aggregation-prone proteins remains unclear. We therefore examined BAG2 condensate behavior under IFNγ in a model of protein aggregation.

### I-PDBs mediate tau fibril capture and clearance at the ER

To test whether the inflammatory remodeling of BAG2 condensates enhances the handling of aggregation-prone substrates, we employed a physiologically relevant model of substrate accumulation and protein misfolding using the tau-P301L mutant, which is associated with frontotemporal dementia. H4 cells were seeded with a synthetic 19aa cognate tau fibril designated jR2R3 P301L which induces stable full-length tau fibrillization that propagates across multiple cell generations^42,43^. Under these conditions, tau was detected both as small, dynamically mobile cytoplasmic fragments and as larger aggregates (Figure 4A, Video S4).

**Figure 4.**
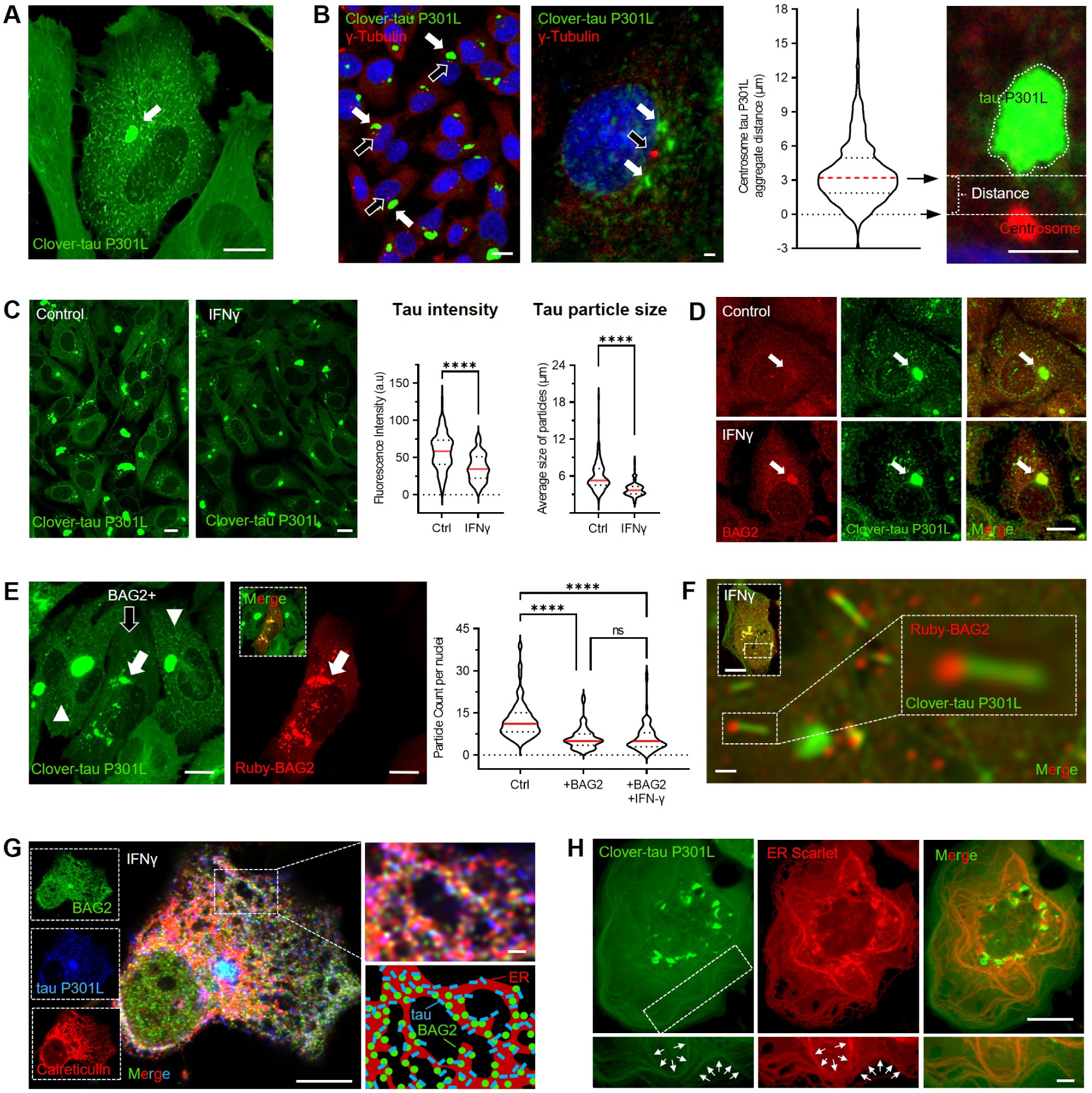
I-PDBs mediate tau fibril capture and clearance at the ER. (A) Live-cell imaging of H4 cells expressing Clover–tau P301L showing intracellular aggregate formation. (B) Immunofluorescence of γ-tubulin (centrosome marker) shows pericentrosomal accumulation of tau aggregates. Open arrows indicate centrosomes; white arrows indicate aggregates. Quantification shows aggregate distance to the centrosome. Violin plots show median (red) and interquartile range (dashed). (C) Clover–tau P301L fluorescence in untreated and IFNγ–treated cells. IFNγ reduces total fluorescence and particle size. Violin plots show median (red) and interquartile range (dashed). Quantification includes 190–270 fields of view from 3 biological replicates per condition. (D) BAG2 immunostaining shows co-localization with tau aggregates in the perinuclear region. (E) Clover–tau P301L cells transiently expressing BAG2. Open arrow indicates a BAG2-positive cell; arrowheads indicate BAG2-negative cells. Quantification of tau particle number per cell in BAG2-positive versus BAG2-negative cells under control and IFNγ conditions. Violin plots show median (red) and interquartile range (dashed). Data from 3 biological replicates. (F) High-magnification image showing co-localization of tau fibrils with ruby-tagged BAG2. (G) Immunofluorescence of BAG2 and calreticulin (ER marker) in tau P301L–expressing cells. Inset and schematic show spatial association of tau, BAG2, and ER membranes. (H) Immunofluorescence of tau P301L–expressing cells with ER-Scarlet. Scale bars, 10 µm and 2 µm (zoom). (C) two-tailed Student’s t test; (E) two-way ANOVA; ****p < 0.0001; ns, not significant. See also Figure S4 and Videos S4-6.

First, to explore the cellular response underlying these phenotypic changes (tau aggregates), we performed bulk RNA-seq analysis comparing H4 cells with and without tau fibrils (Methods). Notably, BAG2 expression was reduced in fibril-bearing cells, which was also confirmed by qPCR (Figure S4B). This was accompanied by transcriptional remodeling indicative of a centrosome/aggresome-directed pathway (Figure S4C). Specifically, dynein–dynactin components (*DYNC1H1, DCTN1-3, ACTR1B*), the centrosomal adaptor *NINL* and the recycling endosome regulator *RAB11FIP3* and motor adaptor *BICD1* were enriched, consistent with enhanced retrograde trafficking toward the microtubule-organizing center. This shift caused by the presence of fibrils is further supported by elevated levels of *HDAC6* and *SQSTM1*, key mediators of aggresome formation, together with enrichment of pericentriolar components *PCM1* and *PCNT*, and reduced factors supporting anterograde transport and microtubule plus-end dynamics (*KIF5B-C* and *MAPRE1-2*) (Figure S4C). Together, these transcriptional changes suggest that tau fibrillization promotes dynein-dependent transport of proteotoxic cargo toward centrosomal aggresome structures.

Large tau aggregates appeared persistently close to the perinuclear region, consistent with the location of aggresome–centrosome structures (Figure 4A). Immunofluorescence analysis using γ-tubulin as a centrosomal marker revealed that large tau aggregates consistently localized in proximity to the centrosome (average distance ∼3 µm) (Figure 4B). This spatial organization highlights a specialized pericentrosomal compartment where aggregation-prone substrates converge under conditions of proteostatic stress^13,42^.

Given the role of I-PDBs in immune-related proteostasis and peptide handling, we next asked whether inflammatory signaling influence tau aggregates. IFNγ caused a marked reduction in total tau fluorescence intensity and average particle size (Figure 4C). To determine whether this response involved BAG2, we first examined the spatial relationship between tau and BAG2 by immunofluorescence. Large tau aggregates showed robust colocalization with endogenous BAG2 following IFNγ (Figure 4D), whereas only weak association was observed under basal conditions. Consistently, overexpression of Ruby–BAG2 confirmed colocalization between BAG2 and tau aggregates (Figure 4E – white arrow) and coincided with a marked reduction in detectable tau species (black arrow vs arrowhead), including reduction in dynamically mobile cytoplasmic fragments (Video S4). Quantitative particle analysis revealed that BAG2 overexpression reduced tau particle number (Figure 4E), with no additional effect of IFNγ, suggesting that BAG2 is rate-limiting for clearance and that IFNγ may act primarily by expanding BAG2 levels and condensation as shown previously (Figure 1A,B). High-resolution imaging further demonstrated that small tau aggregates physically associated with I-PDBs (Figure 4F; Video S4), supporting a role for these structures in sequestering misfolded tau and facilitating its turnover.

To define the subcellular context of BAG2–tau interactions at the ER, we performed immunofluorescence for BAG2 and the ER marker calreticulin in Clover-tau P301L fibril–bearing cells under IFNγ. Endogenous BAG2–ER assemblies (calreticulin) remained closely associated with small tau fibrils (Figure 4G). To better illustrate these interactions, the bottom right inset provides a schematic representation of BAG2–tau contacts at ER-proximal interfaces. Notably, we also observed tight spatial coupling between the ER and the microtubule network even without IFNγ (Figure 4H). Live-cell imaging further revealed that I-PDBs dynamically traverse the ER network, with transient ER rearrangements opening ahead of the moving condensates, illustrating how these assemblies navigate through ER-associated spaces (Video S5). Particularly we can observe the coordinated association among microtubules, small tau aggregates and ER (Video S6), providing a structural framework for the organization and trafficking of target substrates.

ER-associated I-PDBs coordinate with the ER and microtubule network to triage misfolded proteins for degradation, generate antigenic peptides, or sequester material in aggresome-like structures for rerouting to alternative proteostasis pathways^44^. In this way, I-PDBs couple IFNγ-driven proteostasis to peptide production.

### BAG2 couples IFNγ–induced proteostasis to efficient tau peptide production

To establish I-PDBs as immune-associated proteostasis hubs regulating not only fibrillar species but also full-length tau, we examined IFNγ responses in H4 and BAG2 KD cells stably expressing full-length tau. In control cells, western blot analysis revealed a marked reduction in both total and phosphorylated tau following IFNγ (Figure 5A). In contrast, IFNγ had no detectable effect on tau levels in BAG2 KD cells, indicating that BAG2 is required for IFNγ–dependent tau clearance. In addition, whole-cell proteomics in the tau-enriched system showed a similar pattern, with IFNγ producing minimal additional changes in tau abundance in the absence of BAG2 (Figure 5B; Table S1). The tau-enriched background did not alter IFNγ-induced upregulation of BAG2, MHC-I, or PSMB8 (Figure S4D) consistent with our previous findings (Figure 1B, S1C,D). BAG2 KD also did not affect MHC-I levels and IFN-induced MHC-I upregulation (Table S1).

**Figure 5.**
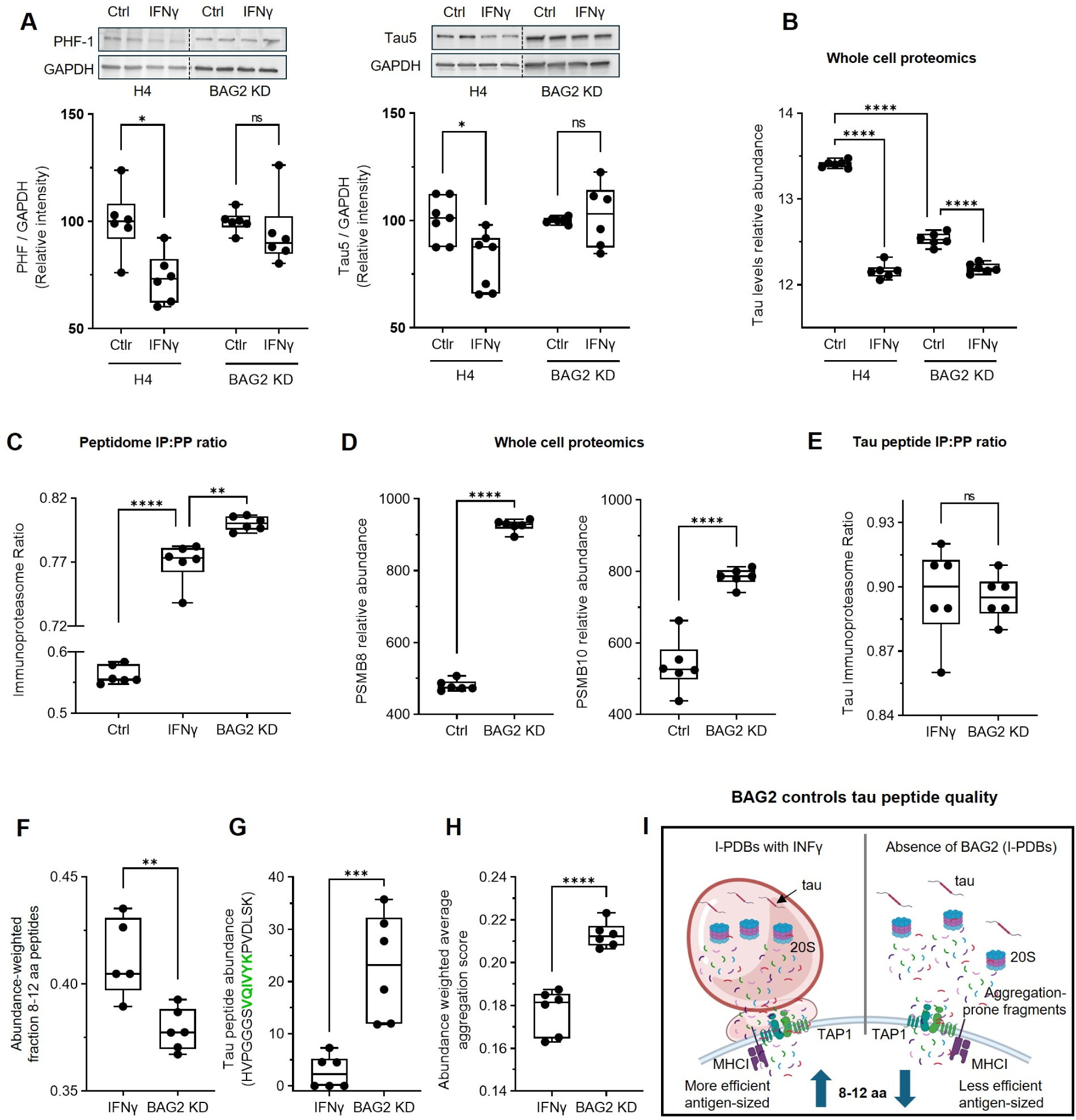
BAG2 couples IFNγ–induced proteostasis to efficient tau peptide production. (A) Immunoblot and quantification of phosphorylated tau (PHF-1) and total tau (Tau5) in H4 and BAG2 KD cells ± IFNγ. IFNγ–induced tau reduction is attenuated in BAG2 KD cells. (B) Whole-cell proteomics of relative tau abundance under the indicated conditions. (C) Ratio of immunoproteasome- to total proteasome-derived endogenous peptides (IP:PP) in untreated, IFNγ–treated, and BAG2 KD tau-overexpressing cells. (D) Proteomics quantification of PSMB8 and PSMB10 abundance. (E) Ratio of immunoproteasome- to total proteasome-derived tau peptides (tau IP:PP) in IFNγ–treated H4 and BAG2 KD cells. (F) Abundance-weighted fraction of immunoproteasome-derived tau peptides of 8–12 aa. (G) Abundance of the PHF6-containing tau peptide (VQIVYK). (H) Aggregation-prone score (7–15-mers) for peptides identified under the indicated conditions. (I) Model: IFNγ induces immunoproteasome remodeling and BAG2 condensate formation at the ER, coupling tau fibril capture to 20S-dependent processing and promoting antigen-sized (8–12 aa) peptide generation; this process is impaired in BAG2-KD cells. Box plots show median (center line), interquartile range (box), and min–max (whiskers); dots indicate independent biological replicates. Statistics: (A–C) two-way ANOVA with Tukey’s test; (D–G) two-tailed Student’s t test. *p < 0.05, **p < 0.01, ***p < 0.001, ****p < 0.0001; ns, not significant. See also Figure S4.

To examine BAG2’s role in processing tau-derived peptides, we conducted an independent peptidome analysis in our tau full-length-enriched system. When calculated across the full peptide set, IFNγ increased immunoproteasome-to-proteasome (IP:PP) ratio (Figure 5C). Notably, BAG2 depletion alone also shifted proteasome output toward the immunoproteasome even exceeding the effect of IFNγ alone. To investigate the mechanism underlying this increase in IP:PP ratio upon BAG2 KD, whole cell proteomics revealed a marked increase in PSMB8 and PSMB10 expression levels (Figure 5D), accompanied by increased levels of autophagy-related markers (Figure S4E).

We next restricted our analysis to tau-derived peptides. Tau peptides were detected primarily in IFNγ-treated cells or in BAG2-deficient cells, whereas in other conditions tau peptides were largely undetectable due to rapid degradation. In both IFNγ and BAG2 KD settings, tau-derived peptides exhibited a strong immunoproteasome bias (∼90%), exceeding that observed across the full peptide set (78%) (Figure 5E), indicating that BAG2 loss, even in the absence of IFNγ, promotes a high immunoproteasome dependence of tau processing. Strikingly, BAG2 depletion reduced the abundance-weighted fraction of tau peptides within the optimal 8–12 aa range compared with IFNγ treatment (Figure 5F). These findings suggest that BAG2 contributes to the efficient generation of appropriately sized tau peptides. Notably, targeted analysis identified increased abundance of a tau-derived peptide encompassing the pathogenic PHF6 motif (HVPGGGS**VQIVYK**PVDLSK) in BAG2-depleted cells compared to IFNγ (Figure 5G).

Parallel analyses of peptide properties showed that BAG2 depletion shifts the overall peptide landscape toward increased amyloidogenicity, as estimated by amyloid-predict scoring (Figure 5H). A schematic model summarizing how I-PDBs control tau peptide quality is presented in Figure 5I. Together, these results indicate that I-PDBs not only facilitate the clearance of full-length tau proteins but also shape the quality of the tau peptide output, functionally linking proteostasis to immune-related processing. We therefore next examined whether I-PDBs are required for antigen-dependent CD8⁺ T cell conjugation.

### I-PDBs are required for CD8⁺ T cell–mediated immune surveillance

To assess the contribution of BAG2 and fibril formation to antigen-dependent CD8⁺ T cell conjugation, we performed co-culture experiments with CD8⁺ T cells and H4 cells. MHC-I typing confirmed compatibility, permitting antigen-specific recognition (Methods). Activated CD8⁺ T cells (Figure S5A,B) were pretreated with IFNγ for 24 h, and then co-cultured with H4 cells (5:1 ratio) for 1h, and conjugation was assessed by immunofluorescence (Figure 6A) and quantified (Figure 6B). IFNγ markedly increased the number of CD8⁺ T cell conjugates compared with untreated H4 cells (Figure 6A,B), indicating that immune activation enhances target cell recognition. In the inset, CD8⁺ T cells extend multiple processes while scanning for antigen consistent with dynamic immune surveillance behavior. In BAG2 KD cells, a pronounced reduction in CD8⁺ T cell conjugation relative to BAG2-expressing cells was observed, despite IFNγ (Figure 6A,B), indicating that BAG2 condensates are required for immune recognition. Notably, approximately 10% of H4 cells escaped CRISPRi-mediated BAG2 depletion (Figure 6A inset, 6C), and CD8⁺ T cells preferentially formed conjugates with this BAG2-positive subpopulation: nearly 51% of conjugates formed with these cells, corresponding to a fivefold BAG2-dependent conjugation index (Figure 6D).

**Figure 6.**
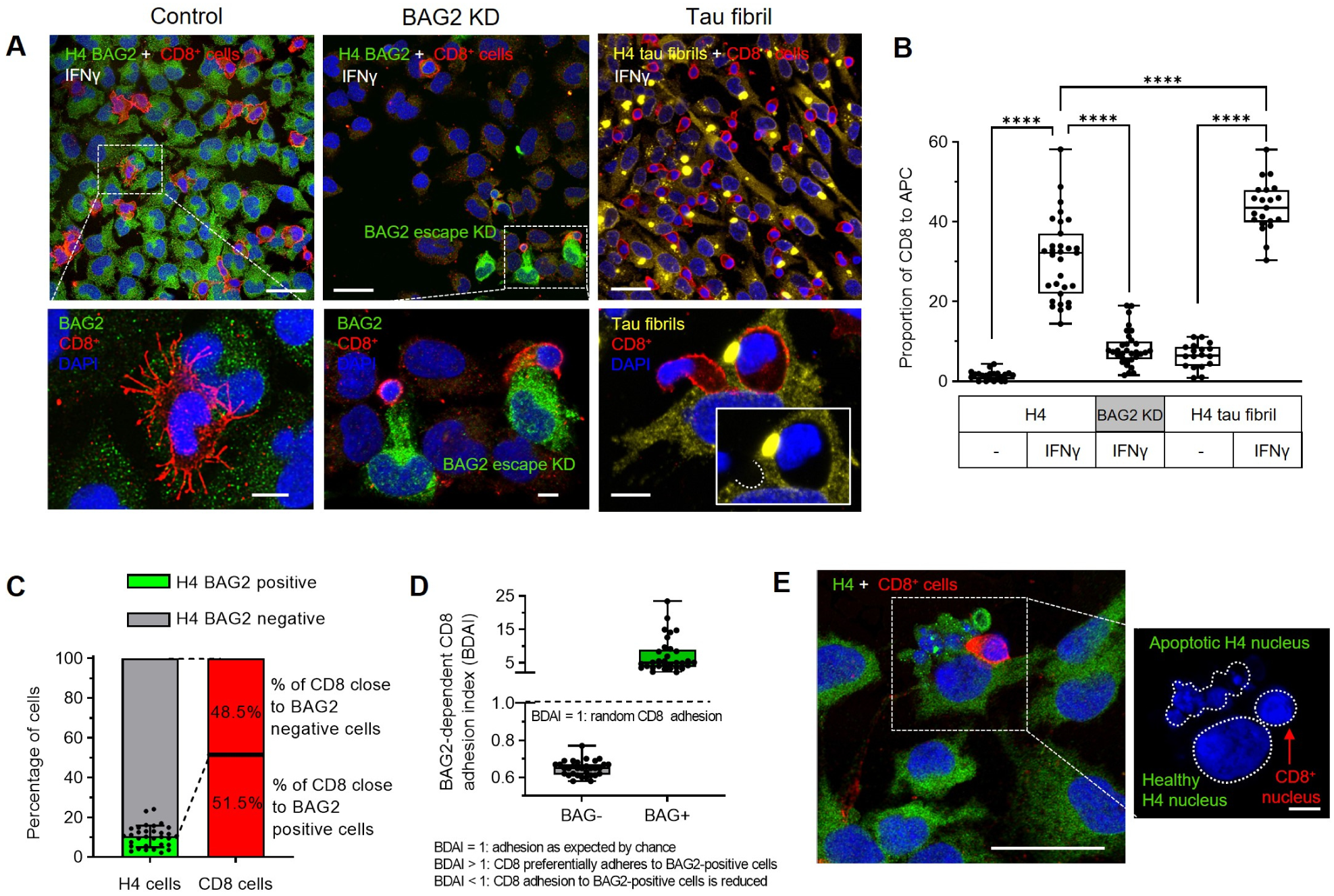
I-PDBs are required for CD8⁺ T cell–mediated immune surveillance. (A) Representative immunofluorescence images of CD8⁺ T cells co-cultured with IFNγ–pretreated H4, BAG2 KD, or tau fibril–containing H4 cells. CD8⁺ T cells were labeled with anti-CD8; BAG2 was detected by immunofluorescence; Clover–tau fibrils were imaged directly. (B) Quantification of CD8⁺ T cells relative to antigen-presenting cells under untreated or IFNγ–treated conditions. (C) Fraction of BAG2-positive and -negative cells within BAG2-KD cultures and corresponding CD8⁺ T cell engagement. (D) BAG2-dependent CD8 adhesion index, reflecting stable T cell attachment. (E) Representative images after 8 h co-culture showing apoptotic morphology in target cells following CD8⁺ T cell interaction. Box plots show median (center line), interquartile range (box), and min–max (whiskers); Each dot represents proportion of H4 to CD8 cells per field of view (FOV) taken from 6 independent biological replicates/condition with an average of 90 H4 cells per FOV in each condition. Statistics: two-way ANOVA with Tukey’s test. ****p < 0.0001. Scale bars, 30 µm and 5 µm (zoom). See also Figure S5.

To evaluate the impact of proteotoxic stress on immune recognition, H4 cells bearing tau-P301L fibrils were co-cultured with CD8⁺ T cells. IFNγ pre-treatment significantly increased CD8⁺ T cell conjugation to fibril-bearing cells compared with their untreated counterparts (Figure 6A,B), indicating that immune activation facilitates recognition of cells with accumulated misfolded proteins. Close contacts between CD8⁺ T cells and H4 cells were frequently observed, with CD8^+^ T cells indenting the target cell cytoplasm without membrane penetration (Figure 6A, right bottom panel), consistent with stabilized engagement.

In extended co-culture experiments (8 h), BAG2-positive H4 cells in proximity to CD8⁺ T cells displayed morphological features consistent with apoptosis, including cell shrinkage, membrane blebbing, and subsequent fragmentation of both cytoplasm and nucleus (Figure 6E; Figure S5C). The spatial association with CD8⁺ T cells supports a cytotoxic, likely granule-mediated mechanism.

Collectively, these findings reveal a cellular strategy for converting intracellular stress into immune signals via the PAIR pathway, establishing I-PDBs as critical hubs with potential relevance across multiple pathological contexts, including neurodegeneration.

## DISCUSSION

Cells must maintain proteome integrity while remaining responsive to immune surveillance. Here we identify a mechanism that links proteotoxicity to immune signaling through BAG2-driven LLPS. In so doing, we define a highly motile membraneless organelle—the Immune-Protein Degradation Bodies (I-PDBs)—characterized by its association with the immunoproteasome located on the cytoplasmic side of the ER where it facilitates trafficking of immunogenic peptides into the ER lumen and ultimately to the MHC-I complex without triggering ERAD or UPR. I-PDBs recruit ISGylation machinery^41^, a pathway linked to antigen presentation^45^, IFNγ signaling components^46^, and reduced ubiquitination^47^. I-PDBs are enriched in the 20S proteasome core which is known to degrade a substantial fraction of the proteome, preferentially targeting intrinsically disordered proteins^48,49^. Within the context of inflammation, I-PDBs are routed along a pathway we define as the PAIR (Proteostasis-Associated Immune Relay) pathway. I-PDBs function on the PAIR pathway to capture misfolded substrates such as tau at points of intercalation between the ER and the microtubules where a process of proteolysis begins that channels the resulting peptides into the ER lumen and toward MHC-I antigen presentation.

We propose that BAG2 acts as the primary scaffold for I-PDBs assembly. Its oligomeric architecture enables multivalent interactions that drive phase separation and client recruitment under stress^13,27^ while phosphorylation further modulates these properties^13,50,51^. These features likely allow BAG2 condensates to function as sweepers that selectively capture proteins and spatially organize proteolysis and antigenic peptides generation. Unlike membrane-bound vesicles, I-PDBs can assemble and disassemble throughout the cytoplasm. Their low energetic barrier for molecular entry and exit enables the selective sequestration of misfolded proteins in ways that vesicular compartments cannot. Moreover, continuous exchange with the surrounding cytoplasm allows I-PDBs to dynamically sample and reflect the proteostatic state of the cell.

Aggregating proteins challenge cellular degradation, as oligomers and fibrils often exceed proteasome and autophagy capacity, accumulating in aggresomes^42^. Interestingly, BAG2 condensates handle misfolded with two different types of condensates depending on the cellular conditions. BAG2 condensates undergo a fundamental reorganization of their constituents and localization from stress BAG2 condensates (S-PDBs) to I-PDBs in the presence of IFNγ. S-PDBs, previously described and designated^13,14^ as BAG2 condensates, assemble under stress conditions such as sucrose, and transit misfolded proteins such as tau to the aggresome. The centrosome, itself a phase-separated structure^52^, often serves as a convergence point for misfolded proteins transported along microtubules to aggresomes^53,54^ which sequester proteins that escape canonical proteasomal degradation.

IFNγ shifts BAG2 condensates from an association with the centrosome to the ER where they are enriched in antigen-processing machinery. By retaining proteins in compartments coupled to antigen processing, I-PDBs appear to enhance peptide generation for MHC-I and promote immune recognition, whereas aggresome sequestration favors autophagic clearance and reduces immunological recognition. Without BAG2, misfolded proteins are routed to aggresome–autophagy–lysosomal pathways.

Immunoproteasome-mediated proteolysis generates a diverse peptide repertoire, including spliced epitopes that expand antigen presentation^55–57^ and bioactive peptides with innate immune functions^58^. This biochemical complexity suggests that spatial organization within I-PDBs may shape the peptide repertoire available for immune recognition. While total peptide levels are reduced with the presence I-PDBs, those peptides of optimal length for efficient antigen presentation compatible with MHC-I motifs^16^ are enriched. Rather than indiscriminately accelerating degradation, BAG2 condensates refine proteolysis. Efficient antigen presentation requires not only the generation of peptides but also preservation from further degradation that can escape immune recognition. Many proteasome-derived fragments are rapidly destroyed by cytosolic peptidases before reaching MHC-I^59,60^ so only a small subset are ultimately presented. Our data suggests that I-PDBs conduct this immune shaping.

Tau protein exemplifies the distinct roles of BAG2 condensates in their different guises. I-PDBs recognize and intercept smaller tau species on the ER–microtubule network before they undergo self-assembly. Notably, in our tau fibril model, I-PDBs retained the capacity to process tau even as larger aggregates accumulate suggesting that PAIR and autophagy pathways operate in parallel. Indeed, large tau aggregates, in the same cell model, show pronounced colocalization with SQSTM1 and VCP^42^, supporting their ongoing engagement with autophagy-dependent pathways. Loss of BAG2 increased immunoproteasome-derived tau peptides (∼90%, versus ∼78%) while reducing the fraction of peptides within the optimal MHC-I length range. In parallel, BAG2 depletion increased the abundance of aggregation-prone fragments containing the PHF6 motif (VQIVYK) and shifted the overall tau-derived peptide repertoire toward increased amyloidogenic signatures. Thus, I-PDBs shape antigen-compatible peptides while limiting aggregation-prone intermediates, ensuring efficient tau processing before irreversible deposits form. These results indicate that immunoproteasome-mediated immune recognition is most effective in the context of I-PDBs.

Loss of BAG2 reduces T cell recognition after IFNγ treatment suggesting that I-PDBs enhance antigen-presenting capacity. Although H4 cells and CD8⁺ T cells are not fully HLA-matched and baseline alloreactivity cannot be excluded, the reduced T cell recognition in absence of BAG2 support I-PDBs actively shaping antigen presentation. Tau fibrils alone did not strongly stimulate CD8⁺ T cell engagement but acted synergistically with IFNγ, exceeding the levels observed in IFNγ-treated H4 cells without tau fibrils. Inflammatory signaling and proteotoxic stress appear to cooperate to expand the pool of substrates available for antigen processing, potentially exacerbating a CD8⁺ T cell response. Our CD8⁺ T cells were not primed against specific tau epitopes, suggesting that these effects reflect a broader remodeling of the cellular immunopeptidome. Although some of the enhancement could reflect increased alloreactivity due to IFNγ-induced MHC-I upregulation, the additional effect of tau fibrils suggests a direct role for proteotoxic stress in modulating T cell engagement.

Cytotoxic CD8⁺ T cells accumulate in brain regions affected by tauopathy^34,38^, raising the possibility that the PAIR pathway modulates neuronal immune recognition. The components underlying the PAIR pathway are broadly expressed across cell types, suggesting that this mechanism is not restricted to one specific cell type. Instead, PAIR may represent a conserved strategy through which cells translate potential proteotoxicity into immune surveillance signals. In this context, tau antigens released from neurons may be taken up by microglia or astrocytes within the CNS, while antigen drainage to peripheral lymphoid organs enables uptake by dendritic cells and subsequent priming of tau-specific CD8⁺ T cells^37^, a process to which the PAIR pathway may also contribute.

By spatially organizing proteolysis within phase-separated compartments, BAG2 condensates couple protein quality control to antigen presentation, enabling cells to communicate their internal proteostasis state to the immune system. More broadly, these findings reveal a spatially organized proteostasis–immunity axis that may operate across diverse physiological and pathological contexts. Whether recruitment contributes to disease progression or serves to prevent tau spread by eliminating toxic cells^61^ suggests the necessity for a balance in the PAIR pathway between beneficial and deleterious effects.

### Limitations of the study

A limitation of this study is the reliance on cellular models, which may not fully capture the complexity of organismal physiology. However, this reductionist approach enabled an exceptional level of mechanistic and structural resolution, integrating multiple complementary methodologies and high-resolution imaging of the intracellular environment that would be difficult to achieve in more complex systems. Importantly, the core components underlying PAIR—such as BAG2, the immunoproteasome, MHC-I, and inflammatory signaling pathways—are broadly conserved, supporting the generalizability of our findings for both normal and pathological processes. Thus, the use of cellular systems is not only justified but was essential to uncover the fundamental organizational principles described here. Future studies extending these observations into more complex *in vivo* models will be important to further validate and expand the physiological relevance of this framework.

## RESOURCE AVAILABILITY

### Lead contact

Further information and requests for resources and reagents should be directed to and will be fulfilled by the lead contact, Daniel C. Carrettiero.

### Materials availability

All unique/stable reagents generated in this study are available from the lead contact upon reasonable request.

### Data and code availability

- The mass spectrometry proteomics data have been deposited to the ProteomeXchange Consortium via the PRIDE partner repository, dataset identifier PXD076373. The bulk RNA-Seq data have been deposited at GEO, accession number GSE326781. Data are currently under controlled access and will be made publicly available upon publication following peer review.
- Original western blot images have been deposited at Mendeley at DOI 10.17632/nbmxvy7g99.1.
- Microscopy data reported in this paper will be shared by the lead contact upon request.
- Original code has been deposited at https://github.com/samlobe/BAG2_condensate_signature and is publicly available at DOI: 10.5281/zenodo.19433054 https://github.com/samlobe/structural-hierarchical-embed and is publicly available at DOI: 10.5281/zenodo.19433066 https://github.com/samlobe/amyloid-predict and is publicly available at DOI: 10.5281/zenodo.18882434
- Any additional information required to reanalyze the data reported in this paper is available from the lead contact upon request.

## ACKNOWLEDGEMENTS

This work was supported by the São Paulo Research Foundation (FAPESP; 2023/05075-0 to D.C.C.) and the National Institutes of Health (NIH; 2R01 AG056058 to K.S.K.). We acknowledge the use of the NRI-MCDB Microscopy Facility, and the Resonant Scanning Confocal microscope supported by NSF MRI grant 1625770. We thank Timothy Tran, Catherine Pugsley, and Mariana G. Arsky for technical support, and Emidio Beraldo-Neto for technical contributions during the early stages of the proteomics work.

## AUTHOR CONTRIBUTION

Author contributions: D.C. Carrettiero, K.S. Kosik, and M.C. Almeida conceived the project. D.C. Carrettiero, M.C. Almeida, A.P. Longhini, C.M. Camargo, E.D. Tinkle, G.Z. Duarte, I. Hirsch, C.A.J. Ribeiro, M. Kwon, and F.A. Oliveira performed the molecular and cellular experiments and analyzed the data. D.C. Carrettiero, M.C. Almeida, and T. Wang performed the proteomics and peptidomics experiments under the supervision of J.A. Steen. S. Lobo performed aggregation propensity analyses and protein structural modeling under the supervision of M.S. Shell and J.-E. Shea. K.S. Kosik and D.C. Carrettiero supervised the study. D.C. Carrettiero, K.S. Kosik, and M.C. Almeida wrote the manuscript. All authors reviewed and edited the manuscript.

## DECLARATION OF GENERATIVE AI AND AI-ASSISTED TECHNOLOGIES

During the preparation of this work, the authors used ChatGPT (OpenAI) to assist with language editing, enhance clarity and text concision. After using this tool, the authors reviewed and edited the content as needed and take full responsibility for the content of the published article.

## METHODS

### EXPERIMENTAL MODEL DETAILS

#### Cell culture and generation of stable cell lines

H4i cells (HLA typing: *A03:01, A30:02; B08:01, B18:01; C05:01, C07:01*), which constitutively express CRISPRi machinery (dCas9–KRAB) together with a TagBFP reporter^62^, as well as BAG2 knockdown (BAG2 KD) cells, were cultured in Dulbecco’s Modified Eagle Medium (DMEM) supplemented with 10% fetal bovine serum (FBS) and 1% penicillin/streptomycin. Cells were maintained at 37 °C in a humidified incubator with 5% CO₂ and were routinely tested negative for mycoplasma contamination.

Single-guide RNA (sgRNA) constructs for dCas9–KRAB–mediated gene repression were cloned into the pLG15 vector as previously described to generate lentiviral particles. These CRISPRi H4i cells (BAG2 KD) were previously generated and characterized by our group^13^. Briefly, H4i cells were transduced with lentivirus encoding either a BAG2-targeting sgRNA (5′-GTGGGGAGCGCAAGTCTCTG-3′) or a non-targeting control sgRNA (scramble), followed by selection with puromycin (1 µg ml⁻¹).

Stable H4 and BAG2 KD cells expressing ruby-tau were generated using a plasmid encoding a neomycin (G418/Geneticin) resistance cassette in cis. Cells were transfected using Lipofectamine 3000 (Invitrogen, Cat#L3000008) according to the manufacturer’s instructions. Twenty-four hours post-transfection, Geneticin (Thermofisher, Cat# 10131035) was added to the culture medium at a final concentration of 500 µg ml⁻¹. Medium was replaced every 2 days, and cells were monitored daily for cytotoxicity. Cells that had stably integrated the plasmid typically survived antibiotic selection for approximately 9 days. Surviving cells were expanded for at least 4 weeks in T75 flasks and subsequently used as polyclonal stable cell lines.

Stable H4 cell lines expressing mClover3-0N4R-P301L tau under a TET-Off expression system were generated by lentiviral transduction of H4 cells. Positive cells were isolated by sorting in purity mode on a Sony MA900 flow cytometer. Expression of mClover3-0N4R-P301L tau was maintained in the absence of tetracycline/doxycycline. These cells are referred to here as tau P301L. To induce templated aggregation, stable mClover3-0N4R-P301L tau-expressing cells were seeded with recombinant jR2R3-P301L tau fibrils as previously described^34^. Briefly, the jR2R3-P301L tau fragment was assembled into fibrils in vitro and sonicated immediately before use to generate short, seeding-competent species. Cells were exposed to 1 µM jR2R3 fibrils delivered with 1.25 µL Lipofectamine 2000 (Thermo Fisher Scientific, Cat# 11668500) per 100 µL media to facilitate cytosolic entry. After 24–48 h, cells were washed and maintained in complete medium. Following seeding, cells were monitored by live-cell fluorescence microscopy for the appearance of discrete mClover3-positive puncta, indicative of intracellular tau aggregation. Cells displaying robust aggregates were isolated under sterile conditions and expanded. Once established, tau aggregates were stably propagated across cell divisions, with daughter cells inheriting mClover3-0N4R-P301L-positive inclusions over multiple passages, consistent with templated self-propagation. These aggregate-propagating lines were used in all subsequent experiments requiring stable intracellular tau fibrils, and are referred to here as tau fibril cells.

Human CD8⁺ T cells (Stem Cell Technologies, Cat#200-0164, Lot#2504418023, HLA typing: *A02:05:01G, A23:01:01G; B40:02:01G, B41:01:01G; C02:02:02G, C07:01:01G*) were thawed and resuspended in RPMI 1640 supplemented with 10% fetal bovine serum (FBS) and 1% penicillin/streptomycin. Cells were counted and adjusted to a density of 1 × 10⁶ cells ml⁻¹. T cells were activated using Dynabeads Human T-Activator CD3/CD28 (Thermo Fisher Scientific, Cat#11131D) at a 1:1 bead-to-cell ratio. Prior to use, beads were washed once in RPMI according to the manufacturer’s instructions.

Cells were cultured at 37 °C in a humidified atmosphere with 5% CO₂ in RPMI supplemented with recombinant human IL-2 (30,000 U ml⁻¹, Milteny Biotec, Cat #130-097-743). IL-2–containing medium was replaced daily. Cells were expanded for up to 8 days, after which CD3/CD28 beads were removed using a magnetic separator. Cells were subsequently rested for approximately 14 h in fresh IL-2–containing medium prior to downstream use or cryopreservation.

For freezing, cells were collected by centrifugation (300 × g, 3 min), resuspended in freezing medium (90% FBS, 10% DMSO), and cryopreserved using a controlled-rate freezing container prior to long-term storage in liquid nitrogen.

### METHOD DETAILS

#### Transient DNA transfection

Cells were transiently transfected using Lipofectamine 3000 according to the manufacturer’s instructions. Briefly, plasmid DNA was diluted in Opti-MEM and combined with P3000 reagent, while Lipofectamine 3000 was diluted separately in Opti-MEM. The DNA–lipid complexes were formed by incubation for 10–15 min at room temperature and then added dropwise to cells at ∼70–80% confluency. Cells were incubated under standard culture conditions and analyzed 24–48 h post-transfection. The following plasmids were used: ER-mScarlet (Addgene plasmid #137805), mClover2-BAG2, and mRuby2-PSMB8. The mClover2-BAG2 and mRuby2-PSMB8 constructs were generated by cloning into the Clover-mRuby2-FRET-10 backbone (Addgene plasmid #58169).

#### APEX2-based proximity labeling proteomics

Proximity labeling was carried out using APEX2 following protocols previously applied to liquid–liquid phase separated condensates^63,64^. Cells were transfected with an APEX2–BAG2–Clover construct, which was generated by cloning into the Clover-mRuby2-FRET-10 backbone (Addgene plasmid #58169). BAG2 served as the proximity labeling bait and mClover2 was used to identify transfected cells. Stable cell populations were generated by Geneticin (G418) selection and further enriched for Clover-positive cells by fluorescence-activated cell sorting using a Sony MA900 cytometer to obtain a homogeneous population.

For proximity labeling, cells were left untreated or pre-treated for 24 h with IFNγ (50ng/ml), followed by incubation with biotin phenol (2.5 mM) in complete growth medium for 3 h at 37°C. Where indicated, sucrose (125 mM) was added during the final hour of incubation in previously untreated cells. Biotinylation was initiated by rapid replacement of the medium with PBS containing Mg²⁺ and Ca²⁺ (PBS++) supplemented with freshly prepared H₂O₂ (100 mM) and allowed to proceed for 60 s. Reactions were quenched immediately using ice-cold stop buffer containing sodium azide, sodium ascorbate, and Trolox, followed by extensive washing. Biotin phenol and H₂O₂ concentrations and reaction times were empirically optimized by titration (**Figure S1E**).

Cells were lysed in RIPA buffer supplemented with protease inhibitors and quenching reagents. Lysates were clarified by centrifugation, protein concentrations were determined, and samples were normalized prior to enrichment. Biotinylated proteins were reduced, alkylated, and enriched using streptavidin-conjugated magnetic beads under denaturing conditions. Beads were thoroughly washed, and on-bead digestion was carried out using sequencing-grade trypsin. The resulting peptides were recovered, quantified using the Thermo Fisher Peptide Assay (colorimetric; cat. #23275), and processed for LC-MS/MS analysis.

#### Sample Preparation and Fractionation Strategy for Proteome and Peptidome Profiling

Cells were subjected to six experimental conditions: untreated H4 cells; IFNγ stimulation (50 ng/mL); BAG2 knockdown (BAG2 KD); BAG2 KD combined with IFNγ; treatment with the immunoproteasome inhibitor KZR-616 (DOSAGE); and KZR-616 combined with IFNγ. Following treatment, cells were incubated for 24 h. Following treatment, cells were washed twice with PBS and rapidly quenched with cold methanol (–20 °C) to preserve protein integrity. The cell layer was scraped, and the suspension was homogenized by ten passages through an 18-gauge needle, followed by ten passages through a finer 25-gauge needle to ensure efficient lysis and DNA shearing. This sequential mechanical disruption reduces sample viscosity, enhances protein solubilization, and improves extraction of both soluble proteins and low–molecular-weight endogenous peptides. Samples were centrifuged to separate the methanol-soluble fraction (supernatant A) from precipitated material. The pellet was resuspended in water, homogenized using the same needle sequence, and centrifuged again to recover re-solubilized proteins (supernatant B). Supernatants A and B were combined to maximize protein recovery. This methanol–aqueous extraction strategy enables recovery of both soluble proteins and a subset of initially precipitated proteins, broadening proteome coverage while preserving endogenous peptides. Combined extracts were dried prior to downstream processing.

For parental H4 cells, dried extracts (A+B) were reconstituted in 1% formic acid (FA) and centrifuged to separate acid-soluble and acid-insoluble fractions. The acid-soluble supernatant, enriched in endogenous peptides, was directly subjected to LC–MS/MS analysis, generating the peptidome dataset. To enable parallel proteome and peptidome analyses, samples were processed using complementary workflows. For Tau H4 cells, the combined dried extract (A+B) was solubilized in a urea-based buffer and subjected to tryptic digestion, generating the whole cell proteome dataset. In parallel, an aliquot of the same extract was treated with FA and centrifuged, and the resulting acid-soluble supernatant was analyzed to generate a matched peptidome dataset under comparable extraction conditions.

#### LC-MS/MS analysis

Evotips were activated through washing with 0.1% formic acid (FA) in Acetonitrile (ACN), incubation in 1-propanol, and washing with 0.1% FA in water. 400 ng of each sample was diluted in 20 μL MS buffer A (0.1% FA in water) and loaded onto Evotips for desalting. Loaded samples were then further washed with MS buffer A prior to LC-MS injection.

LC-MS/MS analysis was performed with an EvoSep One LC system coupled to a timsTOF HT (Bruker). Separation was done using a 30 sample per day (SPD) method, corresponding to a 44-min LC gradient (48-min cycle time) with a 15 cm length × 150 µm I.D. C18 column (Pepsep, Bruker). MS/MS analysis was performed using a data-independent acquisition with parallel accumulation-serial fragmentation (dia-PASEF) mode. Positive polarity MS spectra were collected from 100 to 1700 m/z. MS1 TIMS was set with a 1/K0 range from 0.7 to 1.3 V*s/cm^2^ with a ramp time of 30 ms. MS/MS spectra were collected from 350.7-1250.6 m/z in 60 m/z fixed windows with a 1/K0 range from 0.6-1.6 V*s/cm^2^ using a collision energy calculated on the slope from 20-59 eV on the mobility range.

DIA data were processed in real time using dia-PASEF Spectronaut ® 20 (version 20.2) by Bruker ProteoScape™ (BPS version 2026). Raw files were searched against the reviewed human proteome (UniProt UP000005640, 20,405 entries) with the following modifications: carbamidomethyl Cys (fixed), acetyl (protein N-term, variable), oxidized Met (variable). Digestion was set to fully tryptic with up to two miscleavages. Precursor and protein identifications were filtered at 1% false discovery rate (FDR) using a decoy database strategy, and precursor tolerance was set to 20 ppm. The quantitative peptide and protein summaries were finally generated by Label-free dia-PASEF quant workflow from Spectronaut® version 20.2 IDs (dia-PASEF-LFQ Spectronaut® 20 Quant).

#### Immunofluorescence

After treatments, cell cultures were fixed in a 1:1 methanol-acetone for 10 min at −20°C, washed (3×1 min) with PBS, and permeabilized with 0.1% Triton X for 15 min. Cells were washed 2 × in PBS and incubated for 30 min in blocking buffer at RT (2% NGS, normal goat serum, 4% BSA, 0.2% Triton X-100). Cells were incubated with primary antibody in blocking medium overnight at 4°C, washed (3×5 min) with PBS and incubated for an additional 1 h with secondary antibodies. After washing (3×5 min), the wells were covered in antifade mounting media (ProLong™ Diamond Antifade Mountant with DAPI, Thermofisher, Cat #P36962) and visualized by a Leica SP8 fluorescence microscopy. Primary antibodies used included: rabbit-anti BAG2 (1:300, AbCam, Cat# ab79406); mouse-anti PSMB8 (1:200, Invitrogen, Cat#MA5-15890)mouse-anti calreticulin (1:1000, AbCam, Cat# ab22683); mouse-anti TAP1 (1:100, Thermofisher, Cat# MA5-20071); mouse-anti gamma tubulin (TU30)(1:500, Novus, Cat# NB500-574); mouse-anti human CD8 alpha (1:100; RandD Systems, Cat# MAB3803); mouse anti-MHC class 1 (F-3)(1:200, Santa Cruz Biotechnology, Cat# sc-55582). Secondary antibodies were Goat anti-Rabbit Alexa Fluor Plus 488 (1:400, Thermofisher, Cat#A-11034) and Goat anti-mouse Alexa Fluor 555 (1:400, Thermofisher, Cat# A-21422)

#### Protein extraction and Western blot

Western blotting assay was performed after IFNγ treatment. Briefly, after treatments, cells were lysed with RIPA buffer (1% Triton X-100, 0.5% sodium deoxycholate, 0.1% SDS, 150 mm NaCl, 50 mm TrisHCl, pH 7.4). Protein concentration was estimated by the BCA protein assay kit (PierceBCA Protein Assay, Thermofisher, Cat# 23227) and was adjusted to 1 μg/μl. Total cell lysates (10 – 20 μg) were resolved by SDS-PAGE (Mini-Protean TGX Precast Gels, BioRad, cat# 456-9036) and transferred to a 0.45µm PVDF (Thermofisher,Cat# 88518) or nitrocellulose membrane (GE Healthcare, cat # 10600007). Membranes were incubated in a blocking buffer (PBS, 5% BSA) for 1 h at RT. After overnight incubation (4 C) with primary antibody – rabbit anti-BAG2 1:2000 (AbCam), mouse anti MHC-I, 1:1000 (Santa Cruz Biotechnology), rabbit anti PSMB8, 1:1000 (Invitrogen), rabbit anti-PHF-1, 1:50 (PeterDavis), mouse anti tau5, 1:500 (Thermofisher, Cat#AHB0042) – the blots were washed 2x in Tween 20-TBS (TBS-T) then incubated with IRDye secondary antibodies (1:5000 dilution in Intercept Blocking Buffer, Li-cor, Cat# 927-70001) for 1 h at RT. The blots were then washed 2x in TBS-T and imaged using a digital fluorescent imaging system (ChemiDoc MP Imaging System, BioRad). Quantification of pixel intensity was done with ImageJ and bands were quantified as a percentage of the control. GAPDH (1:5000 Thermofisher, Cat# AM4300) was used as a housekeeping control.

#### RNA isolation, cDNA synthesis, and quantitative PCR

Total RNA, including mRNA, was isolated from cells using the PureLink™ RNA Mini Kit (Thermo Fisher, Cat#12183018A) following the manufacturer’s protocol. Cells were lysed in the provided lysis buffer and homogenized to ensure complete disruption. RNA was purified using silica-based spin columns, with sequential washes to remove contaminants, and eluted in RNase-free water. RNA concentration and purity were measured using a NanoDrop spectrophotometer (Thermo Fisher Scientific). Purified RNA was stored at −80 °C until use.

Complementary DNA (cDNA) was synthesized from equal amounts of total RNA using the High-Capacity cDNA Reverse Transcription Kit (Thermo Fisher, Cat#4368814). Reverse transcription reactions contained reverse transcriptase, reaction buffer, dNTPs, and primers provided with the kit, and were performed according to the manufacturer’s instructions. Synthesized cDNA was diluted as appropriate and either used immediately or stored at −20 °C for downstream analysis.

Quantitative PCR (qPCR) was performed using TaqMan assays (Thermo Fisher Scientific). Reactions were prepared with cDNA, TaqMan Universal PCR Master Mix (Thermofisher, Cat#4444557), and gene-specific probes, and run on a QuantStudio 5 Real-Time PCR System (Thermo Fisher Scientific) under standard cycling conditions. All reactions were performed in technical triplicates. Relative gene expression was calculated using the ΔΔCt method and normalized to housekeeping gene expression.

#### Bulk RNA Sequencing

H4 cells expressing Clover-tau-P301L and either (i) stably harboring tau fibrils (initially seeded with the 4R miniPrion jR2R3 P301L) or (ii) without fibril seeding, were plated in biological quadruplicate (n = 4 per condition), alongside parental H4 cells lacking tau expression as a negative control. Cells were plated in 6-well plates and cultured for 24 h prior to RNA isolation. Total RNA was harvested using TRIzol reagent (Thermo Fisher, Cat#15596026) according to the manufacturer’s instructions.

RNA-seq libraries were prepared from purified total RNA using the TruSeq RNA Sample Preparation Kit (Illumina, Cat#RS-122-2001) following the Low sample protocol. Briefly, poly(A)+ mRNA was purified from total RNA and chemically fragmented prior to first-strand cDNA synthesis, followed by second-strand cDNA synthesis. Double-stranded cDNA underwent end repair, 3′ adenylation, and ligation of indexed adapters. Adapter-ligated fragments were PCR-enriched to generate the final barcoded libraries. Library fragment size distributions were assessed on an Agilent TapeStation, after which libraries were normalized, pooled, and sequenced at the UC Santa Barbara Biological Nanostructures Laboratory (BNL) core on an Illumina NextSeq platform using 75-bp single-end reads.

### QUANTIFICATION AND STATISTICAL ANALYSIS

#### Differential Proteome and Immunopeptidome Analyses

Label-free precursor ion intensities were obtained from mass spectrometry datasets (APEX2-BAG2 proteomics, whole cell proteomics, and immunopeptidome). For all datasets, intensities were log2-transformed (log2(x + 1)). Features with >25% missing values within any condition were excluded, and remaining missing values were imputed using k-nearest neighbors (KNN).

Differential abundance was determined using linear models with empirical Bayes moderation implemented in limma. Pairwise contrasts were defined according to experimental design, and p values were adjusted using the Benjamini–Hochberg method. Features with FDR < 0.05 were considered significantly differentially abundant.

Pathway enrichment analysis of significantly regulated proteins was performed using Enrichr^65^ with Reactome and WikiPathways databases.

#### Quantification of Immunoproteasome-to-Total Proteasome Cleavage Ratios

DIA-MS–derived endogenous peptide data from H4 cells across six experimental conditions were used. Precursor quantities were summed across charge states for each peptide, and C-terminal residues were used to classify peptides as immunoproteasome-derived (F, Y, L, I, V, W, M, R, K) or constitutive proteasome–derived (D, E, S, T, A, G, Q, N). The ratio of immunoproteasome-derived to total proteasome-derived peptides was calculated per sample, and differences between conditions were assessed by two-way ANOVA with treatment and cell type as factors, followed by Tukey’s post hoc test.

#### Immunopeptidome Length Distribution Analysis

Raw precursor ion intensities were extracted from the mass spectrometry output and used as a quantitative measure of peptide abundance. For each biological replicate, peptides were grouped according to sequence length, and precursor intensities corresponding to peptides of 8–12 amino acids were summed to obtain the total 8–12 aa abundance per sample. To account for inter-sample variability in total peptide yield, ionization efficiency, and instrument response, the summed abundance of 8–12 aa peptides was normalized to the total precursor intensity of all identified peptides within the same sample. The relative contribution of 8–12 aa peptides to the peptidome was calculated as: Fraction (8–12 aa) = Σ (precursor intensity of 8–12 aa peptides) / Σ (precursor intensity of all detected peptides).

This fractional abundance metric reflects changes in the proportional representation of 8–12 aa peptides independent of global shifts in total peptide abundance. Statistical comparisons were performed using a two-way analysis of variance (ANOVA) with treatment and cell type as independent factors. When significant main effects or interactions were detected, Tukey’s post hoc multiple comparisons test was applied to adjust for multiple testing. Statistical analyses were conducted in Prism (version 10.1.2), and adjusted p values < 0.05 were considered statistically significant.

#### Protein language modeling analysis

We trained a full-protein classifier to distinguish proteins enriched in the BAG2-APEX IFNγ proteome from proteins detected in the IFNγ whole-cell proteome, using ESM3 sequence+structure embeddings derived from AlphaFold-predicted structures. Positive proteins were defined as proteins detected in the APEX2-BAG2 dataset under IFNγ treatment, and negative proteins were defined as proteins detected in the whole cell proteomics IFNγ dataset but absent from the APEX2-BAG2 IFNγ dataset. Proteins present in both lists were assigned to the positive class. Model performance was evaluated across 20 random protein-level 70/30 train/test splits using a pipeline consisting of L1-based feature selection followed by L2 regularized logistic regression. The median ROC AUC was 0.836, with 16th and 84th percentile values of 0.823 and 0.856, respectively. As a control for abundance-related confounding, we also evaluated baseline protein abundance alone (log10_U), which performed substantially worse (median ROC AUC 0.585). To identify interpretable local signatures, we decomposed protein structures into contiguous secondary-structure composites using DSSP-style 3-state secondary-structure assignments. We then generated cropped-local ESM3 sequence+structure embeddings for local motifs. We focused on solvent-exposed 3-element motifs of length 25-35 aa. We considered motifs with more than 40% solvent-exposed residues; Individual residues were considered solvent-exposed if their relative solvent-accessible surface area exceeded 0.2. Motifs were scored using the trained classifier. High-scoring versus low-scoring motifs were defined as the top and bottom quartiles of motif scores, respectively, and we evaluated interpretable composition features that distinguished high-scoring and low-scoring motifs. For motif-level interpretation, we evaluated a panel of sequence-composition features, including the fraction of each amino acid and grouped residue classes measured in either the full motif sequence or the exposed residues only. Grouped features included acidic (D/E), positive (K/R/H), charged (D/E/K/R and D/E/K/R/H), polar (S/T/N/Q), hydrophobic (A/V/I/L/F/Y/W/M), aromatic (F/Y/W), and amide (N/Q) residues, as well as net charge summaries. P values were computed using two-sided Mann-Whitney U tests and adjusted for multiple hypothesis testing using the Benjamini-Hochberg procedure. Across all solvent-exposed 3-element motifs of length 25-35 aa, the strongest interpretable feature associated with higher classifier scores was the fraction of acidic residues (D/E), which was greater in high-scoring than in low-scoring motifs (median 0.160 vs 0.097; p = 2.0 x 10^-172, q = 1.3 x 10^-170).

#### Aggregation propensity analysis

Aggregation propensity was estimated using the general model from amyloid-predict (https://github.com/samlobe/amyloid-predict). Only peptides 6-15 amino acids in length were included to match the sequence-length range used in amyloid-predict’s training. For each biological replicate, peptide-level aggregation scores were weighted by peptide abundance to calculate an abundance-weighted mean aggregation score. Higher values indicate a greater contribution of peptides with higher predicted aggregation propensity to the overall peptide pool.

#### Bulk RNA Sequencing Analysis

FastQC was used to assess sequencing quality, and all libraries passed standard QC metrics. Adapter sequences were removed from reads prior to quantification. Trimmed reads were quantified against the human reference transcriptome (hg38) using Salmon. Differential expression analysis was performed in R using DESeq2. Variance-stabilizing transformed (VST) counts were used for dimensionality reduction and visualization; UMAP embeddings were generated to assess replicate concordance and sample separation. Replicates clustered tightly by condition, and the experimental groups were well separated in UMAP space.

Differential expression testing was performed in DESeq2 using pairwise contrasts between the experimental conditions. In addition to transcriptome-wide analysis, we specifically interrogated a curated gene set derived from prior proteomic analyses to determine whether these candidates also showed transcriptional changes. For each pairwise comparison, log2 fold changes and associated statistical significance values were extracted from the DESeq2 results for the curated genes and used to assess concordance between the proteomic and transcriptomic datasets.

#### Image analysis

Fluorescence imaging (in vivo and ex vivo) was performed using a resonant scanning confocal microscope (Leica SP8), with laser lines sequentially acquired in line-scan mode to minimize spectral overlap. Live-cell imaging was conducted in a temperature- and CO₂-controlled environmental chamber maintained at 37 °C and 5% CO₂. Image visualization and analysis were performed using Leica Application Suite X and Imaris (v11.0.1; Oxford Instruments), unless otherwise specified. Quantification of condensate number and size (Figure 1), distances between condensates (Figure 2), and distances from the centrosome to large tau aggregates (Figure 4) were carried out in Imaris. Objects below 0.20 µm—approximating the lateral resolution limit of confocal microscopy—were excluded from analysis. Image deconvolution was performed to improve contrast and resolution by reducing out-of-focus signal. Deconvolution was carried out using Huygens Essential (v18.10; Scientific Volume Imaging) with the classical maximum likelihood estimation (CMLE) algorithm, using a signal-to-noise ratio of 20 and 40 iterations

For Clover-tau P301L fluorescence intensity, particle size and count per nuclei, image analysis was done in the ImageJ (Fiji) software v2.14.0/1.54r. tau intensity was quantified by thresholding the tau channel and measuring the integrated density (IntDen) in the region of interest (ROI) of the field of view (FOV). Size and number of tau particles were calculated using the analyze particles function on Fiji. The tau channel was thresholded to select all particles in the FOV, then the average size of particles per FOV was calculated. For the BAG2 transfected cells, first the BAG2 channel was thresholded and applied as a mask in the tau channel, and only particles in BAG2-positive or BAG2-negative cells were selected for analysis.

### Statistical analysis

All statistical analyses were performed using GraphPad Prism (v9). Statistical significance was assessed using one-way ANOVA followed by Dunnett’s or Tukey’s multiple comparisons tests, two-way ANOVA followed by Tukey’s multiple comparisons test, or unpaired two-tailed Student’s t-test, as appropriate. A p value < 0.05 was considered statistically significant. Additional statistical analyses are specified in the corresponding figure legends or Method Details.

