## Supplementary material for "BAG2 Condensates Couple Proteostasis to CD8^+^T Cell Surveillance": Figures S1,S2,S3,S4 and S5

### SUPPLEMENTAL INFORMATION

**Table S1**. Mass spectrometry and Bulk RNA-Sequencing analysis results.

**Video S1**. Live-cell imaging of H4 cells co-expressing Clover–BAG2 and Ruby–PSMB8 revealed partial dynamic demixing. Blend mode in Imaris software (Opacity applied to voxels – 3D impression).

**Video S2**. Live-cell imaging of H4 cells co-expressing ER–mScarlet and Clover–BAG2 under IFNγ revealed that I-PDBs localize along ER-associated structures.

**Video S3**. Live-cell imaging of H4 cells co-expressing ER–mScarlet and Clover–BAG2 under sucrose stress revealed the emergence of BAG2 condensates in the cytoplasm, followed by their migration toward the ER.

**Video S4**. Live-cell imaging of a stable H4 Clover–tau P301L fibrillization model revealed small, dynamically mobile cytoplasmic fragments as well as larger aggregates. Co-expression of Ruby–BAG2 confirmed co-localization of BAG2 with tau aggregates.

**Video S5**. Live-cell imaging of H4 cells co-expressing ER–mScarlet and Clover–BAG2 revealed that I-PDBs dynamically traverse the ER network under IFNγ treatment.

**Video S6.** Live-cell imaging of a stable H4 cell model expressing Clover–tau P301L and transiently transfected ER–mScarlet revealed associations among microtubules, small tau aggregates, and the ER.

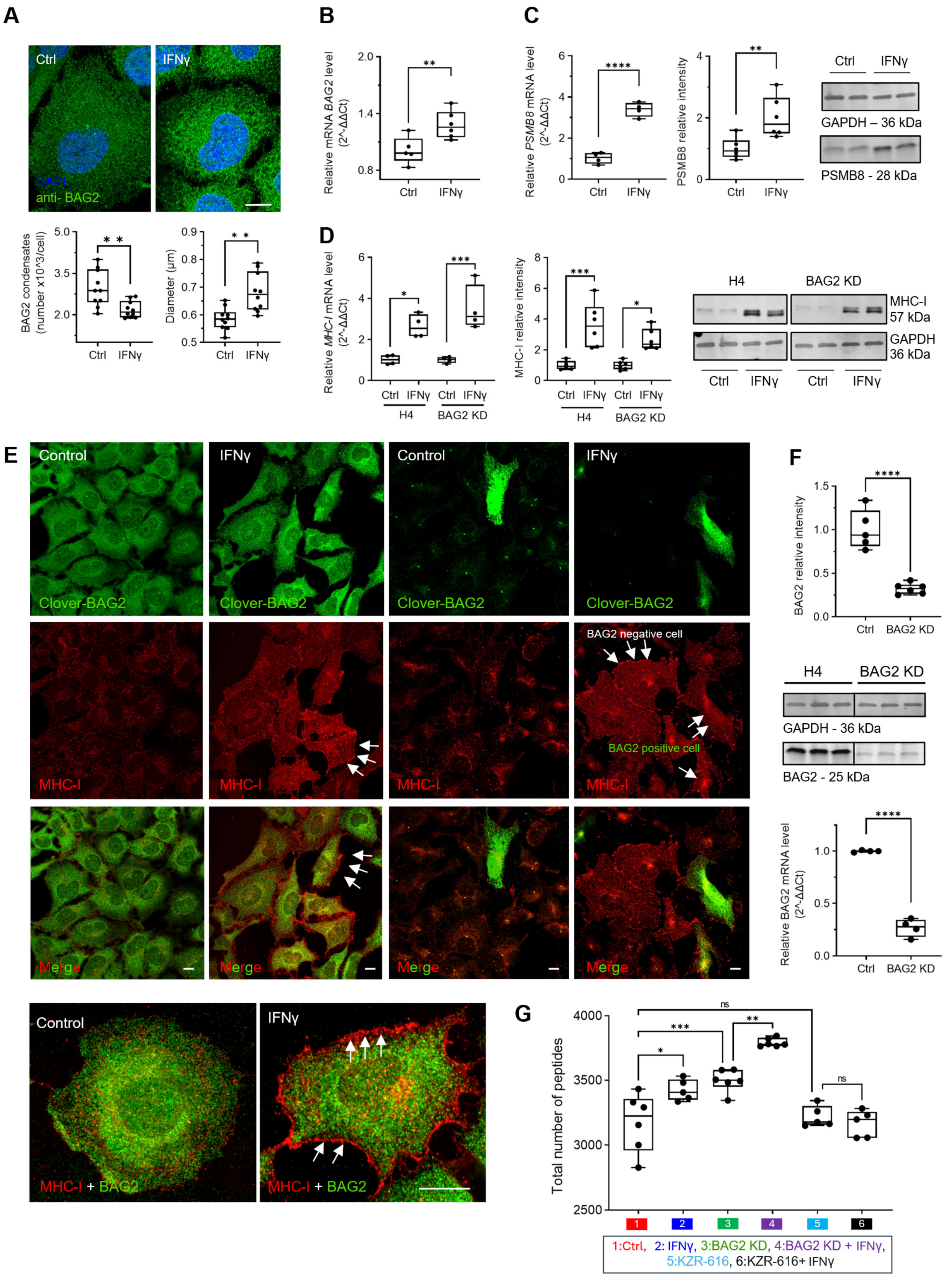
A
Ctrl
IFNγ
DAPI
anti- BAG2
BAG2 condensates (number x10^3/cell)
Diameter (μm)
B
Relative mRNA BAG2 level (2^-ΔΔCt)
C
Relative PSMB8 mRNA level (2^-ΔΔCt)
PSMB8 relative intensity
GAPDH – 36 kDa
PSMB8 - 28 kDa
D
Relative MHC-I mRNA level (2^-ΔΔCt)
MHC-I relative intensity
H4
BAG2 KD
MHC-I 57 kDa
GAPDH 36 kDa
E
Control
Clover-BAG2
MHC-I
Merge
BAG2 negative cell
BAG2 positive cell
F
BAG2 relative intensity
GAPDH - 36 kDa
BAG2 - 25 kDa
Relative BAG2 mRNA level (2^-ΔΔCt)
MHC-I + BAG2
G
Total number of peptides
1:Ctrl, 2: IFNγ, 3:BAG2 KD, 4:BAG2 KD + IFNγ,
5:KZR-616, 6:KZR-616+ IFNγ

**Figure S1. Characterization of BAG2, immunoproteasome components, and MHC-I across IFNγ and BAG2 knockdown conditions.**
(A) Immunofluorescence imaging (top) of endogenous BAG2 in untreated or IFNγ–treated cells (50 ng/mL, 24 h), with quantification (bottom) of condensates per cell. Scale bar: 10 μm. Dots represent mean values per field of view from independent replicates.
(B) *BAG2* mRNA (qPCR) levels in untreated and IFNγ–treated cells.
(C) *PSMB8* mRNA (qPCR) and protein levels in control and IFNγ-treated H4 cells. Representative blots are shown. GAPDH was used as a loading control.
(D) *MHC-I* mRNA (qPCR) and protein levels in control and IFNγ-treated H4 or BAG2 KD cells. Representative blots are shown. GAPDH was used as a loading control.
(E) Immunofluorescence analysis of BAG2 (Clover-BAG2) and MHC-I in H4 and H4 BAG2-KD cells under control and IFNγ-treated conditions. Arrows indicate peripheral localization of MHC-I observed in both cell lines. Representative images are shown.
(F) Western blot (top) and qPCR (bottom) for BAG2 in H4 and BAG2 KD cells.
(G) Total number of peptides detected by peptidomics under the indicated conditions: untreated H4, IFNγ-treated H4, untreated BAG2-KD, IFNγ-treated BAG2-KD, KZR-616-treated H4, and KZR-616+ IFNγ-treated H4 cells.
All box plots display the median (center line), interquartile range (box), and minimum–maximum values (whiskers). Each dot represents one biological replicate. Statistical significance was determined using a two-tailed Student's *t*-test (A, B, F) or 2-way ANOVA followed by Tukey's test (C, D, G) (**$p < 0.01$, ***$p < 0.001$, ****$p < 0.0001$). See also Figure 1

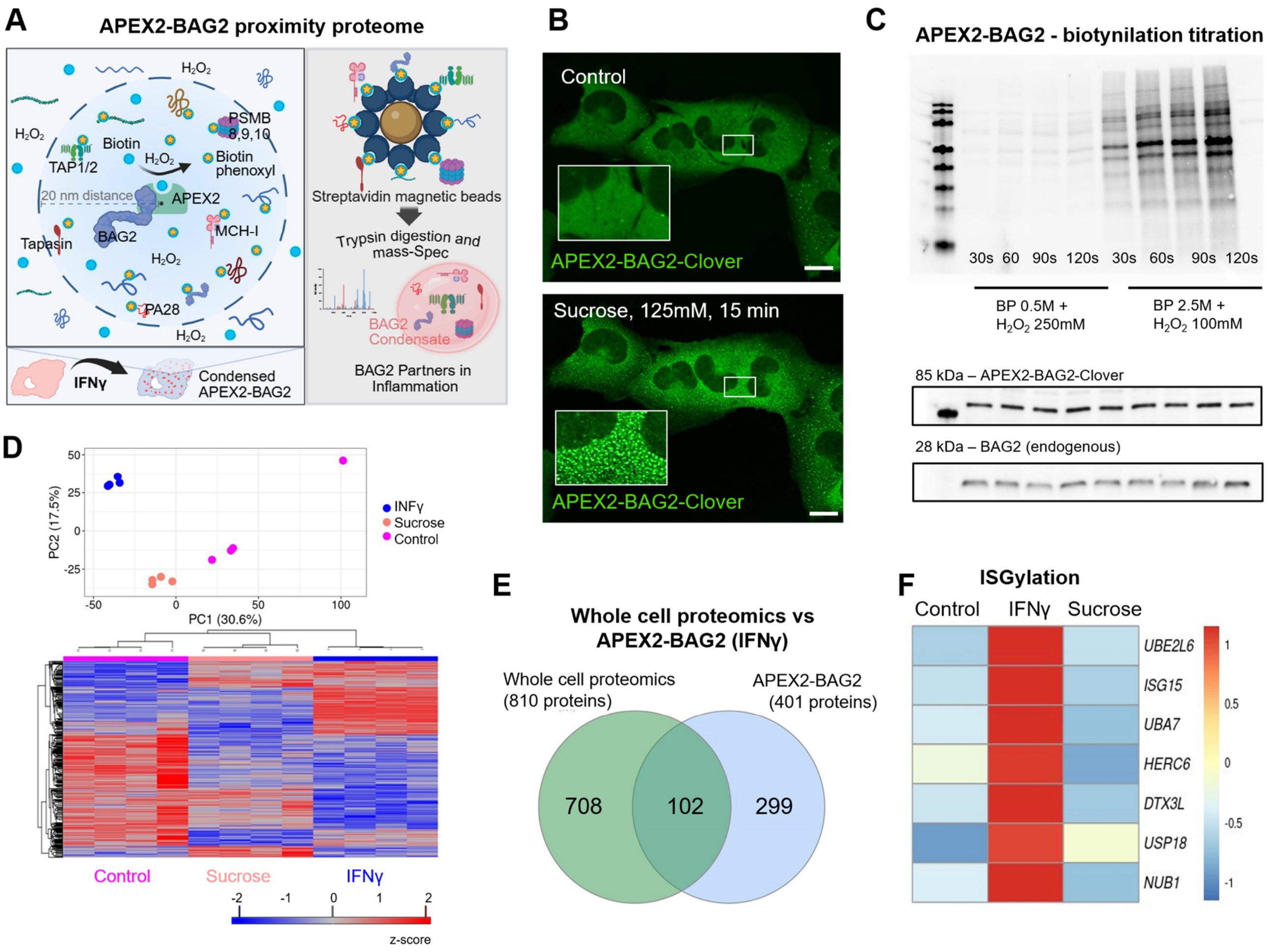


**Figure S2. APEX2-based proximity proteomics defines the BAG2 condensate interactome and its sequence determinants.**

(A) Schematic of the APEX2-BAG2 proximity labeling strategy (Created with BioRender).

(B) Live-cell imaging of cells expressing APEX2–BAG2–Clover before and after osmotic stress induced by sucrose treatment (125mm, 15 min), showing BAG2 condensate formation.

(C) Optimization of APEX2-mediated proximity labeling. Streptavidin–AF680 blot detecting biotinylated proteins following biotinylation titration in APEX2–BAG2 cells. BAG2 immunoblot is shown as an expression control. BP: biotin phenol

(D) Quality control of proximity labeling proteomics. Principal component analysis (top) and heatmap of mean protein abundance (bottom) showing clear separation between treatment conditions.

(E) Overlap between IFNγ–regulated proteins and the APEX2-BAG2 proteome. Whole cell proteomics identified 810 IFNγ-responsive proteins, while APEX2-BAG2 captured 401 proximal proteins, with 299 proteins uniquely identified by APEX2-BAG2, highlighting the specificity of proximity labeling for BAG2 condensates. Notably, shared IFNγ-responsive proteins such as TAP1, PSMB8/9/10, HLA-C, and B2M showed higher log2 fold change in APEX2-BAG2 compared with whole cell proteomics, suggesting a preferential enrichment of these proteins within BAG2 condensates rather than solely reflecting global abundance changes.

(F) Heatmap of proteins associated with the ISGylation pathway identified in the APEX2-BAG2 proximal proteome under untreated, IFNγ-treated, and sucrose-treated conditions. Columns represent the average normalized abundance for each treatment group.

See also Figure 2.

**A**

| Feature | S-PDB (Sucrose-induced) | I-PDB (IFNγ-induced) |
|---|---|---|
| Nature of Stimulus | Physicochemical stress or increased protein crowding/misfolding | Cytokine-driven immune signaling (IFNγ) |
| Cellular purpose | Protect proteome from misfolding and aggregation | Enhanced antigen processing for immune surveillance |
| BAG2 condensate | Rapid driven by client load and molecular crowding | Slower. Transcriptionally dependent |
| BAG2 upregulation | No | Yes |
| Core components | HSP70 and protein quality-control machinery | HSP70 and core Immune machinery |
| Ubiquitin system | No | No |
| 20S engagement | Limited | Strong enrichment |
| ER association | Yes | Yes, stronger |
| Immunoproteasome subunits | No | Yes |
| MHC-I – Peptide loading complex | No | Yes |
| Functional interpretation | General proteostasis condensate supporting stress tolerance | Immune-specialized degradation hub supporting antigen generation |

**B**

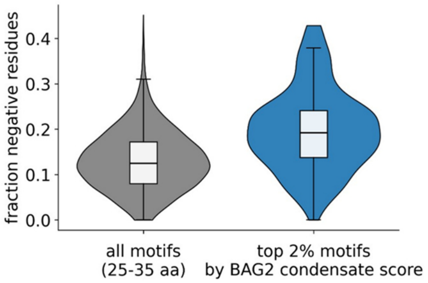


**C**

Calreticulin
disordered-loop-section

B2M
strand-loop-strand motif

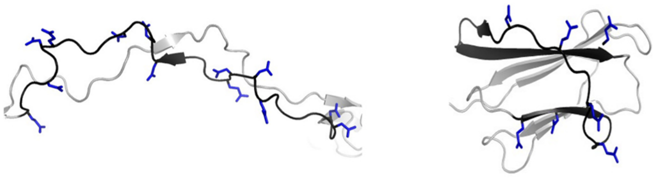

Score: 0.99
Residues 210-240
Sequence:DPDASKPEDWDERAKIDDPTD
SKPEDWDKPE

Score: 0.96
Residues 56-90
Sequence:EVDLLKNGERIEKVEHSDLSF
SKDWSFYLLYYTEF

**Figure S3. BAG2 condensates display distinct features under hyperosmotic and IFNγ stress.**
(A) Summary of shared and distinct features of BAG2 condensates formed in response to sucrose or IFNγ.
(B) Distribution of the fraction of negatively charged residues (D/E) across local 25-35 aa motifs from structured protein regions. Compared with all analyzed motifs, the top 2% highest-scoring motifs by BAG2 condensate score are shifted toward greater acidic residue content, consistent with acidic sequence enrichment as a local signature associated with BAG2 condensate recruitment.
(C) Representative high-scoring local motifs from BAG2 condensate-associated proteins illustrating this acidic sequence signature. Shown are motifs from calreticulin (residues 210-240; 42% negatively charged) and B2M (residues 56-90; 23% negatively charged), both with BAG2 condensate scores in the top 2%. Structures are based on AlphaFold-predicted models. In each structure, the selected motif is highlighted in black and negatively charged side chains (D/E) are shown in blue. See also Figure 2

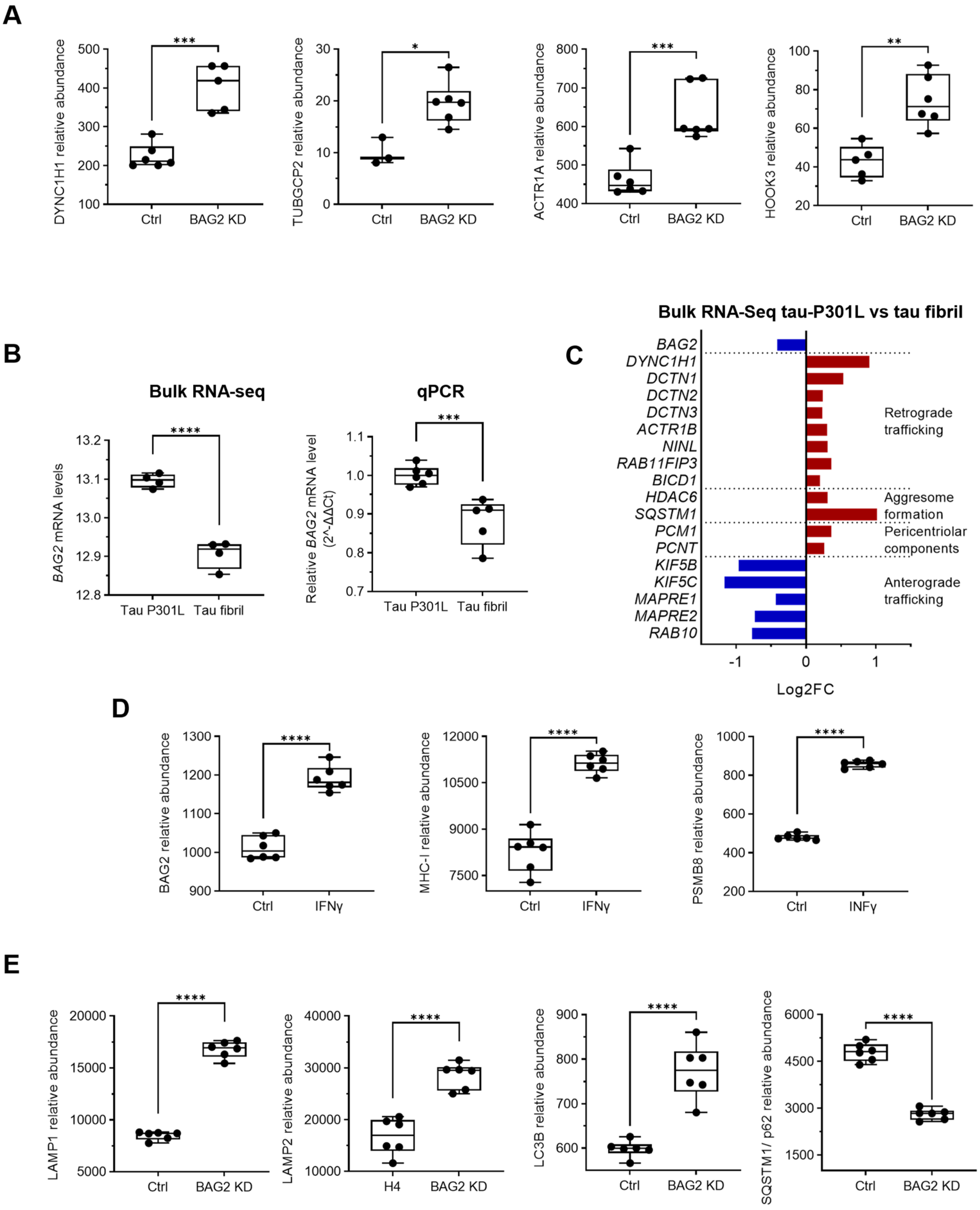
A
DYNC1H1 relative abundance
TUBGCP2 relative abundance
ACTR1A relative abundance
HOOK3 relative abundance
Ctrl
BAG2 KD
***
*
***
**
B
Bulk RNA-seq
qPCR
BAG2 mRNA levels
Relative BAG2 mRNA level (2^-ΔΔCt)
Tau P301L
Tau fibril
****
***
C
Bulk RNA-Seq tau-P301L vs tau fibril
BAG2
DYNC1H1
DCTN1
DCTN2
DCTN3
ACTR1B
NINL
RAB11FIP3
BICD1
HDAC6
SQSTM1
PCM1
PCNT
KIF5B
KIF5C
MAPRE1
MAPRE2
RAB10
Retrograde trafficking
Aggresome formation
Pericentriolar components
Anterograde trafficking
Log2FC
D
BAG2 relative abundance
MHC-I relative abundance
PSMB8 relative abundance
Ctrl
IFNγ
INFγ
****
E
LAMP1 relative abundance
LAMP2 relative abundance
LC3B relative abundance
SQSTM1/ p62 relative abundance
Ctrl
H4
BAG2 KD
****

**Figure S4. BAG2 depletion alters centrosomal trafficking components, gene expression, and autophagy-related pathways.**
(A) Relative abundance of centrosomal and retrograde transport–associated proteins in H4 and BAG2 KD cells, as determined by proteomics analysis.
(B) *BAG2* mRNA levels measured by bulk RNA-seq and qPCR in tau P301L–expressing cells and tau fibril–treated cells.
(C) Divergent plot showing log2 fold change (log2FC) of *BAG2* and trafficking-related genes from bulk RNA-seq analysis in tau fibril–treated cells compared to non-seeded controls. All genes shown have a false discovery rate (FDR) < 0.05. The pattern of gene expression changes is consistent with impaired trafficking and potential cargo accumulation.
(D) Abundance of BAG2, MHC-I and PSMB8 in control and INFγ treated cells, as determined by proteomics analysis.
(E) Abundance of autophagy-related markers in BAG2 KD compared to H4 cells, as determined by proteomics. The combination of decreased SQSTM1 (p62) and increased lysosomal/autophagy markers (e.g., LAMP1, LAMP2, LC3B) is consistent with enhanced autophagic flux, suggesting active cargo degradation rather than accumulation.
Box plots display the median (center line), interquartile range (box), and minimum–maximum values (whiskers). Each dot represents one biological replicate. Statistical significance was assessed using a two-tailed Student's t-test (*p < 0.01, **p < 0.001, ***p < 0.0001, ****p < 0.00001).
See also Figures 3-5.

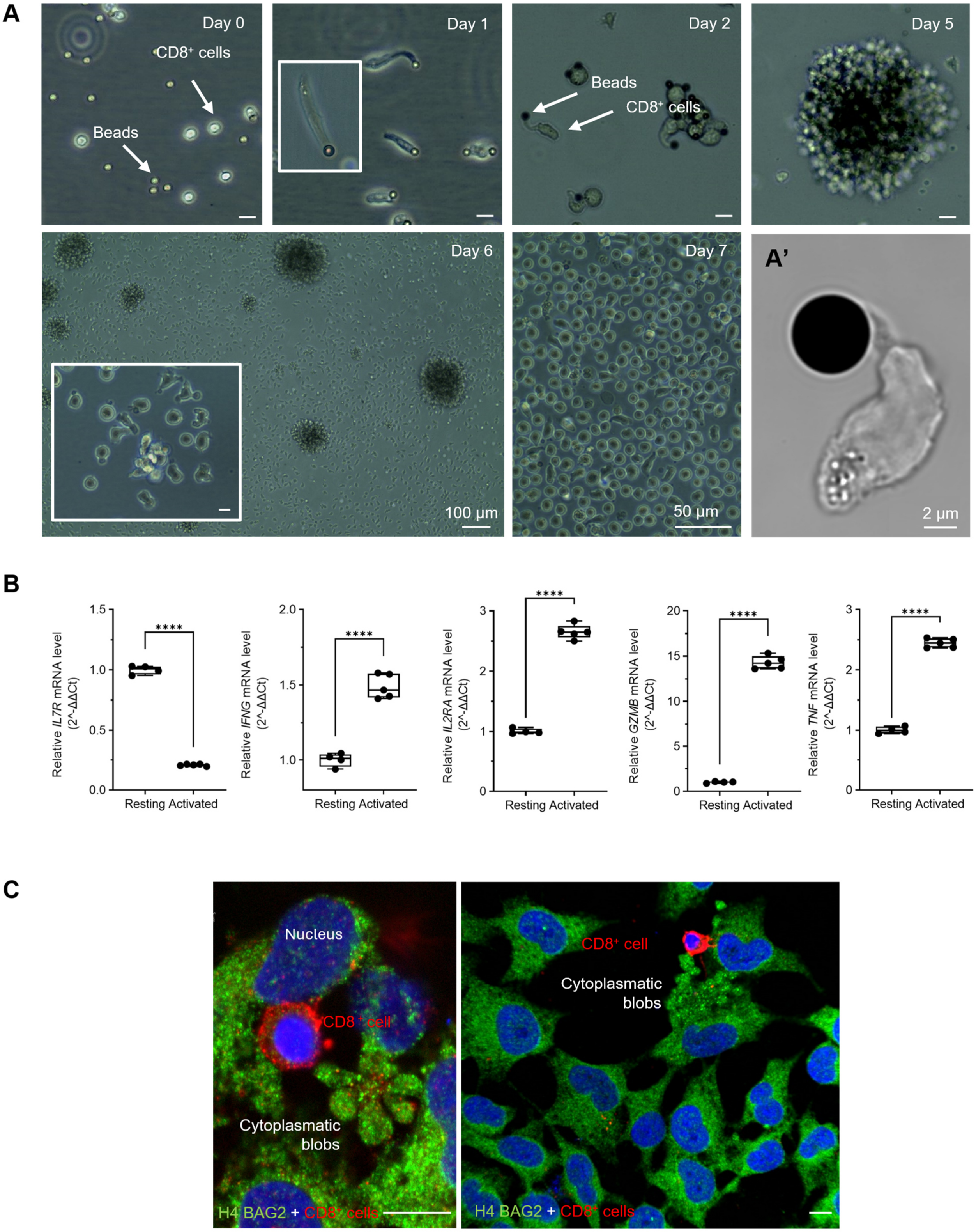
A
Day 0
CD8⁺ cells
Beads
Day 1
Day 2
Beads
CD8⁺ cells
Day 5
Day 6
100 µm
Day 7
50 µm
A'
2 µm
B
Relative IL7R mRNA level (2^-ΔΔCt)
Relative IFNG mRNA level (2^-ΔΔCt)
Relative IL2RA mRNA level (2^-ΔΔCt)
Relative GZMB mRNA level (2^-ΔΔCt)
Relative TNF mRNA level (2^-ΔΔCt)
Resting Activated
****
C
Nucleus
CD8⁺ cell
Cytoplasmatic blobs
H4 BAG2 + CD8⁺ cells
CD8⁺ cell
Cytoplasmatic blobs
H4 BAG2 + CD8⁺ cells

**Figure S5: Activation and expansion of CD8 T cells under dynabead and IL-2 stimulation**

(A) Brightfield images showing CD8 T cell activation and expansion over time in the presence of dynabeads (anti-CD3 and anti-CD28) and IL-2. (A') Confocal brightfield image highlighting the interaction between Ddynabeads and CD8 T cells, as well as morphological changes in T cell, a classical indicator of CD8 T cell activation.

(B) qPCR analysis of activated $CD8^+$ T cells compared with resting CD8 T cells. Box plots represent the median (center line), interquartile range (box), and minimum–maximum values (whiskers). Each dot corresponds to one biological replicate. Statistical significance was assessed using a two-tailed Student's t-test (**$p < 0.01$, ***$p < 0.001$, ****$p < 0.0001$).

(C) Illustrative photomicrographs of H4 cells co cultured with activated $CD8^+$ T cells for 8 h and immunostained for BAG2, illustrating H4 cells with cytoplasmatic blobbing which is a sign of apoptosis.

Scale bar: 10 µm, unless otherwise noted. See also Figure 6.
